# MreB is dispensable for viability but critical for rod shape, motility and biofilm fitness in *Pseudomonas aeruginosa*

**DOI:** 10.64898/2026.08.02.742333

**Authors:** Merve Nur Tunç, Matteo Gérard, Aurélien Barbotin, Marie-Françoise Noirot-Gros, Marina Grégoire, Pierre-Emmanuel Douarre, Arnaud Bridier, Marie Delaby, Yves V. Brun, Steven L. Porter, Romain Briandet, Rut Carballido-López

## Abstract

Despite growing interest in the MreBCD morphogenetic complex as a potential antimicrobial target, its function in *Pseudomonas aeruginosa* remains poorly understood. While previous studies using the MreB inhibitor A22 have established its role in cell shape maintenance and pilus regulation, the impact of *mreB* deletion has not been comprehensively investigated. Using genetic and microscopy-based approaches, we show that deletion of *mreB* is viable in *P. aeruginosa,* but results in spherical cells that lose all forms of motility despite retaining flagella. Importantly, we uncover a previously overlooked polar effect of the in-frame *mreB* deletion on the downstream *mreCD* genes and show, using CRISPRi-mediated silencing, that *mreCD* expression is essential for viability. *ΔmreB* mutants also display increased sensitivity to β-lactam antibiotics and enhanced initial surface attachment, yet form more compact biofilms with reduced dispersal. In mixed-culture biofilms, spherical *ΔmreB* cells are outcompeted by rod-shaped wild-type cells and remain confined to the biofilm base. The identification of natural *P. aeruginosa* isolates carrying truncated *mreB* alleles further indicates that loss of MreB function can be tolerated in natural populations. Together, our findings reveal important contributions of the MreBCD system to viability, morphogenesis, motility and biofilm development in *P. aeruginosa*, providing new insights into bacterial adaptation and informing the development of targeted antimicrobial strategies.

## Introduction

*Pseudomonas aeruginosa* is a highly adaptable, Gram-negative pathogenic bacterium known for its ability to cause a wide range of infections. It is a leading cause of hospital-acquired infections, particularly in immunocompromised individuals, and is frequently associated with respiratory, urinary, wound, and bloodstream infections. Its intrinsic resistance mechanisms and high capacity for acquiring additional resistance genes have made it a prominent member of the ESKAPE group of pathogens, an acronym for the most clinically relevant multidrug-resistant bacteria that effectively “escape” the action of antibiotics. This makes *P. aeruginosa* a major concern in clinical settings and highlights the urgency of studying its resistance mechanisms and treatment strategies^1^. The remarkable ability of *P. aeruginosa* to transition from a single-cell lifestyle to a biofilm-forming community contributes significantly to its resilience in hostile environments, including in the host during antibiotic treatment^2^. This resilience is further reinforced by intrinsic features such as reduced permeability of the double-membrane envelope and highly efficient efflux pumps^3^. Beyond these molecular defences, its characteristic rod-shaped morphology is closely linked to essential cellular processes, such as cell division and motility, which, in turn, support its adaptability and survival strategies in diverse, often hostile environments. Thus, understanding the mechanisms that maintain *P. aeruginosa* rod-shape offers insights into bacterial physiology and potential targets for antimicrobial strategies.

The primary determinant of bacterial cell shape is the peptidoglycan (PG) cell wall, a mesh-like structure that confers rigidity and maintains cellular integrity against internal turgor pressure. PG consists of linear glycan strands composed of alternating N-acetylglucosamine and N-acetylmuramic acid residues, which are cross-linked by short peptide bridges to form a resilient three-dimensional matrix^4^. PG is synthesized by large, dynamic multiprotein complexes that coordinate two enzymatic activities: transglycosylation (TG), which polymerises glycan strands, and transpeptidation (TP), which forms peptide cross-links^5^. In rod-shaped bacteria, two main PG-synthesising machineries operate: the elongasome (also called the Rod complex), associated with the actin-like cytoskeletal protein MreB and responsible for sidewall elongation, and the divisome, organised around the tubulin-like FtsZ and responsible for septum formation during cell division^6,7^. Both complexes rely on SEDS (Shape, Elongation, Division, and Sporulation) proteins for TG activity and on Penicillin Binding Proteins of Class b (bPBPs) for TP activity. Outside the elongasome and the divisome but acting in coordination with them, the bi-functional class A PBPs (aPBPs), which have both TG and TP activity, contribute independently to PG synthesis. In *P. aeruginosa*, aBPs are PBP1a (encoded by *ponA*) and PBP1b (encoded by *mrcB*), and the major bPBPs include PBP2 (encoded by *mrdA*), essential for cell elongation, and PBP3 (encoded by *ftsI*), essential for septum formation^5,8^. Notably, PBP1a has been shown to interact with PBP2 and contributes primarily to elongation, whereas PBP1b has been associated with both elongation and division^9^. Studies in *Escherichia coli* and *Bacillus subtilis* have demonstrated that the Rod complex and aPBPs function in coordinated but largely independent pathways during elongation, underscoring their distinct but complementary roles in maintaining rod shape^10^. In *P. aeruginosa*, the Rod complex specifically relies on the SEDS-family protein RodA (for TG activity) and the bPBP, PBP2 (for TP activity), alongside several accessory proteins that regulate complex stability and localization^5,11,12^. Central to the Rod complex’s function are the actin homolog MreB and its associated partners MreC and MreD, which play critical structural and regulatory roles in the spatiotemporal control of PG insertion during cell growth^10,13^. Through its dynamic polymerisation, MreB is thought to act as a scaffold that limits the diffusion of other elongasome components in the membrane and orients their motion around the cell circumference, thereby promoting cylindrical expansion. In contrast, aPBPs insert PG in an isotropic manner, inserting PG diffusively across the cell surface, whereas the elongasome ensures highly organized circumferential PG insertion in radial hoops^10,12^.

MreB is highly conserved in rod-shaped bacteria, both Gram-positive and Gram-negative, and its depletion leads to severe morphological defects, often resulting in a transition from rod to spherical cell shape, compromised growth rates, and altered cellular function^16^. *mreB* is essential under normal growth conditions in most bacteria in which it has been investigated, including the model Gram-negative and Gram-positive bacteria, respectively, *E. coli* and *B. subtillis*^17,18^. In *P. aeruginosa*, *mreB* was initially proposed to be essential for cell viability when targeted by the antibacterial indole compound CBR-4830, which specifically inhibits MreB, leading to a transition from rod-shaped to coccoid cells and suggesting a critical role of MreB in maintaining bacterial morphology and survival^19^. However, it was recently reported that deletion of *mreB* in *Pseudomonas fluorescens,* a species closely related to *P. aeruginosa,* generates viable spherical cells with reduced fitness, increased cell volume, and perturbed cell division. This finding is consistent with a previous report in which adaptive, viable spherical *mreB* mutants were obtained in *P. fluorescens* via transposon mutagenesis^20^. Additionally, CRISPRi-silenced *mreBCD* cells of *P. fluorescens* resulted in inflated, round cells, with some cells ultimately bursting in overnight culture^21^. In *P. aeruginosa*, Treatment with the MreB inhibitor S-(3,4-dichlorobenzyl) isothiourea (A22)^22,23^ was shown to cause altered pilus production, localization and function in addition to cell rounding^19^. However, it has been suggested that A22 may have additional targets in *P. aeruginosa*^24^, and thus the role of MreB in motility remains to be conclusively demonstrated.

While the role of MreB in cell morphogenesis has been extensively studied, its impact on biofilm formation remains largely unexplored. In *P. aeruginosa*, A22 inhibited biofilm formation and motility of some (but not all) clinical strains at sub-inhibitory concentrations, significantly reducing swarming and twitching motilities, but was not effective in destroying pre-formed biofilms, suggesting that MreB affects the initial stages of adhesion. However, A22 may affect other processes, such as cellular signalling^24^, and thus the direct role of MreB in biofilm formation remains to be demonstrated. Interestingly, in a co-culture biofilm, wild-type *E. coli* cells were predominantly found at the colony base and edges, whereas *E. coli* cells carrying single amino acid mutations in *mreB* that result in altered cell aspect ratio (shorter, wider rods) tended to migrate to the top layers of the biofilm^26^. This finding suggests that cell shape may drive spatial organisation within bacterial communities. The differential positioning occurs as rod-shaped cells form wedge-like structures that burrow beneath spherical cells, enabled by their alignment with surfaces and neighbouring cells. This mechanism, validated through both modelling and experiments with MreB mutants, explains how cell morphology directly influences bacterial positioning and competition within biofilms. However, further research is needed to determine whether similar shape-driven biofilm patterning occurs in *P. aeruginosa* and how it evolves during dynamic biofilm development.

In most rod-shaped bacteria, the two integral membrane proteins MreC and MreD are encoded by the same (*mre*) operon as MreB and are part of the elongasome complex^27^. MreC and MreD are also found in coccoid and ovococcoid species such as *Staphylococcus aureus* and *Streptococcus pneumoniae*, which lack MreB. MreC is a bitopic protein suggested to serve as a periplasmic adaptor, linking MreB to the PG-synthesising enzymes^28^. Additionally, MreC is thought to regulate Rod complex activity by promoting activation of the SEDS transglucosylase RodA and the bPBP transpeptidase PBP2^29,30^. In *P. aeruginosa*, although less well characterized, a recent study suggested that MreC depletion leads to morphological defects and compromised cell envelope integrity^28^. It has also been reported that increased *mreC* expression correlates with enhanced biofilm formation and elevated gentamicin resistance in *P. aeruginos*^31^.

MreD is a polytopic membrane protein believed to help stabilize interactions between MreC and the plasma membrane^32–34^. Both MreC and MreD have been shown to be essential for viability in several rod-shaped bacteria, including *E. coli*^34,35^*, Caulobacter crescentus*, and *B. subtilis*^36^. In these organisms, depletion of *mreC* or *mreD* results in profound morphological defects-cells lose their rod shape, become enlarged or spherical, and ultimately lyse under normal growth conditions. However, in spherical bacteria like *S. aureus*, which lack MreB but retain MreC and MreD, these proteins appear non-essential, and their loss does not significantly affect cell morphology^37^. Similarly, in the ovococcoid bacterium *S. pneumoniae*, MreC and MreD are required for growth only when PBP1a is functional. Their essentiality is alleviated when PBP1a is inactivated, making them dispensable for viability, although some cell division defects may persist^38^. Notably, the essentiality and precise role of MreC and MreD in Pseudomonas species have not yet been thoroughly addressed, representing an important gap in our understanding of elongasome function in these bacteria.

Understanding the role of MreB and its associated proteins in *P. aeruginosa* is critical for elucidating the cytoskeletal underpinnings of bacterial physiology and may offer new avenues for antimicrobial strategies targeting cytoskeletal components. However, the phenotypic consequences of *mreB* deletion in *P. aeruginosa*, particularly in terms of growth mode, potential compensatory genetic adaptations, and biofilm formation, remain poorly characterized. In this study, we investigate the impact of *mreB* deletion in *P. aeruginosa* using a multifaceted approach combining genetic deletions, CRISPR interference (CRISPRi), and live-cell imaging at both the single-cell and biofilm levels. We analysed changes in cellular morphology, division patterns, cell growth kinetics, motility, antibiotic sensitivity and biofilm development. Additionally, we explored whether *mreB* deletion is polar on expression of the downstream *mreCD* genes, and assessed the viability and morphology of MreCD-depleted cells. Our study sheds light on the role of MreB in maintaining structural integrity and adaptive capacity in *P. aeruginosa*.

## Results

### *mreB* null mutants are viable and spherical in *P. aeruginosa*

*In E. coli, mreB* is essential for cell viability in LB medium ^34,35^. In *P. aeruginosa*, it was initially reported to be essential, based on the inability to delete *mreB* unless a second functional copy was present in the cell and the failure to obtain *mreB* insertion mutants in a genome-wide transposon mutagenesis study^19^. However, *mreB* was recently reported to be dispensable for growth in *P. fluorescens*^39^. To determine if *mreB* is essential in *P. aeruginosa,* we attempted to construct in-frame *mreB* deletions in the parental strain PAO1-fix using a conjugation-based method. *mreB* mutants were viable and spherical in LB medium (Fig. 1a and Fig. S1a). Whole-genome sequencing of six independent Δ*mreB* clones revealed varying single-nucleotide polymorphisms (SNPs) in three of them (PAMNT044, PAMNT049, PAMNT051) and insertions in four of them (PAMNT043, PAMNT048, PAMNT049 and PAMNT054) (Table S1). Despite these variations, all six Δ*mreB* mutant strains displayed similar growth rates (Fig. S1b) and spherical morphology (Fig. S1a). We next focused on the analysis of strain PAMNT054 that showed no evidence of suppressor mutations (hereafter called Δ*mreB*). We monitored the growth of wild-type (WT) and Δ*mreB* mutant cells over 140 minutes on 1 % LB agar pads by time-lapse microscopy (Fig. 1a). Because of the clumping of Δ*mreB* cells precluding proper segmentation at later time points, only the first 20 min of these time-lapses were used to quantify cell dimensions. Single-cell growth rate and duplication time were analysed from the first 25 frames (120 min). The average cell length of WT cells was 2.86 ± 0.62 µm, while the length of Δ*mreB* cells was 1.62 ± 0.32 µm (Fig. 1b). While cell length decreases in the absence of *mreB*, the width increased from 0.88 ± 0.08 µm (WT cells) to 1.36 ± 0.19 µm (Fig. 1c).

**Figure 1:**
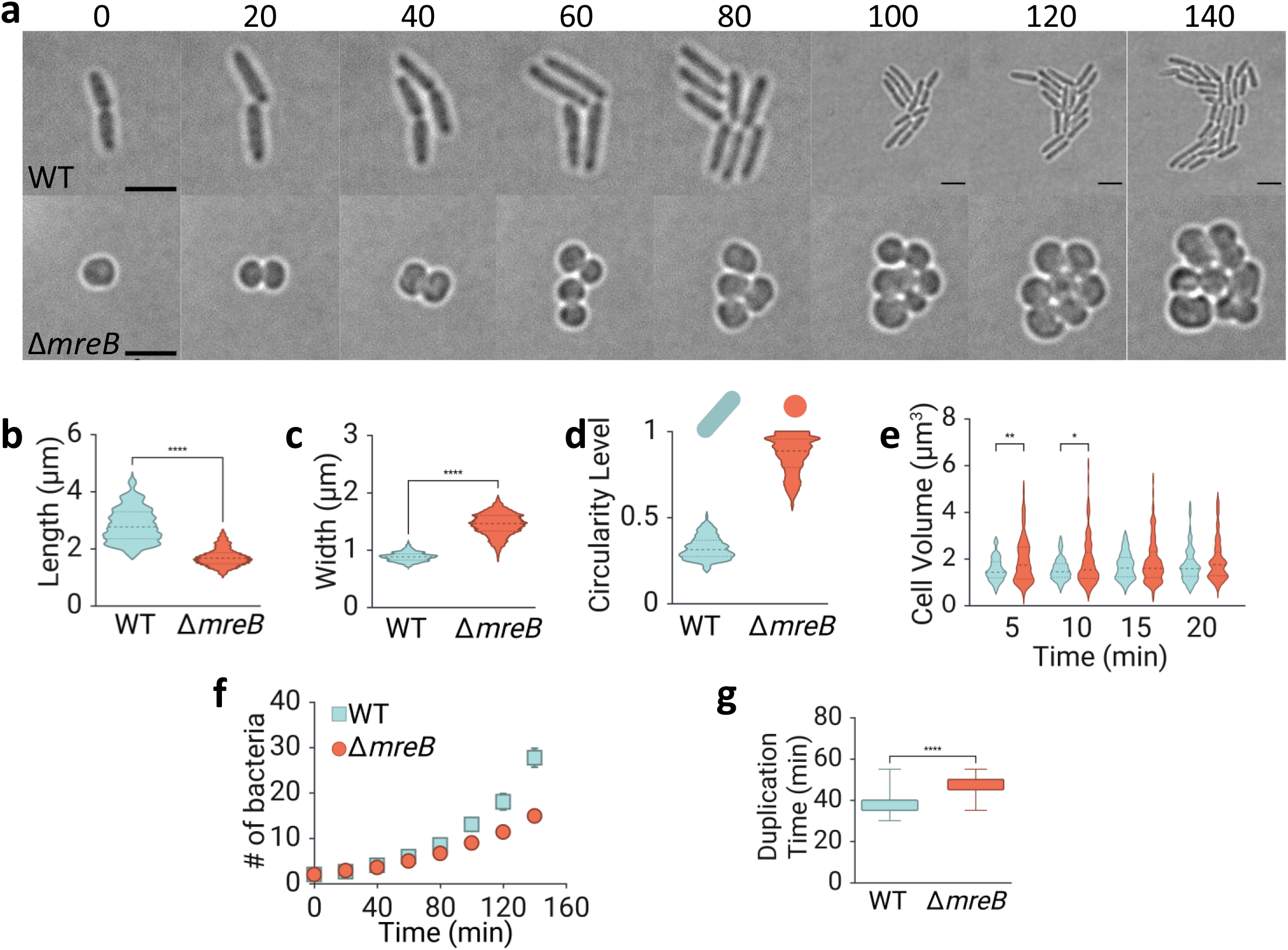
Analysis of the cell structure of *ΔmreB* mutant at the single-cell level. **(a)** Time-lapse bright-field images of *P. aeruginosa* WT PAO1-fix (top row) and Δ*mreB* mutant (bottom row) cells growing on 1% LB agar pads at 37 °C. Images shown correspond to 20-minute over 140 minutes (accompanying movie acquired at 5-minute intervals for 205 minutes. (**S. Movie 1a, b**). Scale bar, 3 μm (**b–e**) Morpho analysis of Δ*mreB* mutant cells (orange) and WT cells (blue) extracted from cells grown on 1% LB agarose pad for 20 min. The first time point was discarded for quantifying cell dimensions to stabilise them on the pad (Fig. S2a, b, d, e). Cell length **(b)**, cell width **(c)**, cell circularity **(d)** and cell volume over time **(e)**. Data are presented as violin plots where the top and bottom lines correspond to the 25th and 75th percentiles, and the middle-dashed line corresponds to the median. Single-cell duplication time **(g)** and number of cells **(f)** of WT and Δ*mreB* strains growing for 120 min on the agarose pad were generated from the same image acquisition used in morphological analysis. Data in **(f)** is presented as a box plot with minimum and maximum data lines. Statistical analyses: Mann–Whitney U test (unpaired, nonparametric, two-tailed) was used for panels **(b)**, **(c)**, **(d)**, and **(g)**; two-way ANOVA with Bonferroni multiple comparisons was used for cell volume over time **(e)**.

WT cells exhibited lower circularity values, reflecting their rod shape. In contrast, Δ*mreB* cells displayed an almost spherical morphology (Fig. 1d). Quantification of cell volume from the first four time points (Fig. 1e) revealed that *ΔmreB* cells had a higher median cell volume than WT cells, but only during the first 10 minutes. Although *ΔmreB* cells with volumes around 5 µm³ were consistently observed at all time points, no WT cells reached this volume range.

Over time, WT cells maintained their characteristic rod shape and continued to divide in an organized manner, resulting in the formation of evenly distributed colonies. In contrast, the *ΔmreB* cells exhibited misoriented division planes and aggregated into irregular, disorganized clusters rather than spreading uniformly. Additionally, *ΔmreB* cells showed reduced population expansion over time, as evidenced by a lower cell count (Fig. 1f). Since no signs of cell lysis were observed, the diminished growth was attributed to a slower growth rate, which was further confirmed by an increased duplication time (47.58 ± 5.44 min) than WT (36.66 ± 1.25 min) (Fig. 1g). Analysis of cell shape and dimensions revealed that *ΔmreB* cells were shorter, wider, and more spherical compared to WT cells. Importantly, the differences in cell volume were time-dependent; *ΔmreB* cells were not consistently larger than WT cells across all time points, highlighting that mreB is dispensable for growth but critical for maintaining rod shape and stable cell dimensions.

### *mreCD* are essential for survival and rod-shaped in *P. aeruginosa*

In-frame and frameshift deletions of *mreB* have been reported to require the expression in *trans* of all three *mreBCD* for complementation of the cell shape defects in both *E coli*^35,40,41^ and *B. subtilis*^17^, suggesting that *mreB* mutants may be polar on expression of the downstream *mreCD* genes. In *E. coli*, *mreC* mutants have been shown to be similarly polar on the expression of *mreD*, and all three *mre* genes are similarly required for rod shape and viability^35,42^. However, the levels of MreC have not been assessed in any Δ*mreB* mutant so far. Western blot analysis using antibodies raised against *P. aeruginosa* MreC showed that our unsuppressed *ΔmreB* strain, as well as the other in-frame deletion mutants isolated in this study, do have reduced levels of MreC relative to the WT strain (Fig. S3a), confirming a polar effect. An extra copy of *mreBCD* was added to the Δ*mreB* and WT backgrounds. While there is no morphological change in WT, Δ*mreB* became rod-shaped again (Fig. S4a, b). We then investigated whether *mreCD* might affect the phenotypes described for the Δ*mreB* mutant. Our attempts to construct a null, in-frame *mreC* mutant using the same chromosomal deletion strategy employed for Δ*mreB* were unsuccessful, suggesting that *mreC* is essential in *P. aeruginosa*. We then used a CRISPR interference (CRISPRi) system to selectively silence either *mreBCD* or *mreCD* in the WT and *ΔmreB* backgrounds. We adapted a CRISPRi system that functions through the coordinated activity of catalytically dead CRISPR-associated protein 9 (dCas9) and a single-guide RNA (sgRNA), previously described in *Neisseria meningitidis* and *P. fluorescens*^21,43^. The system we optimised, gene silencing, is finely tuned via two orthogonal small-molecule inputs: cumate and theophylline in a two-plasmid system. Expression of dcas9 is under control of a cumate-inducible promoter repressed by cumate repressor protein (CymR); in the absence of cumate, CymR binds the (cumate operator) CuO operator to block transcription, while cumate addition relieves this repression, inducing dcas9 (Fig. S5a). In addition, the expression of dCas9 is regulated by a theophylline-responsive OFF riboswitch, which exerts negative control. Upon theophylline binding, a conformational change occurs that sequesters the ribosome-binding site (RBS), thereby inhibiting translation and enhancing repression of dCas9 expression in the absence of cumate. In this dual-control expression system, dCas9 expression is stringently repressed in the presence of theophylline and is induced upon cumate addition. Meanwhile, sgRNA is constitutively expressed. (Fig. S5b). A dCas9-GFP fusion was used as a proxy to assess dCas9 expression at the cellular level. The results showed that expression was homogeneous throughout the population, as revealed by flow cytometry (Fig. S6a), and was modulated by the presence of cumate and theophylline (Fig. S6b). Growth was unaffected by the addition of 200 µM cumate to log-phase and exponentially growing *P. aeruginosa* strains, including WT, *the ΔmreB* mutant, and a HspK-silencing strain used as a control (Fig. S7). HspK is a HyperSpank promoter that has no function in *P. aeruginosa*. Since it will not affect any of the *P. aeruginosa regulations*, we used it as a control to compare the growth rate of WT with the 2-plasmid CRISPRi system. HpsK silenced strain (WT + g^HspK^) strain growing as slowly as the mreB mutant, suggesting that the expression of the CRISPRi plasmids has a small impact on fitness/ growth, as expected from *P. aeruginosa* keeping plasmids (Fig. S7). We next monitored the impact of silencing the *mreBCD* and *mreCD* genes on growth. The *ΔmreB* mutant displayed a slower growth rate and reached a lower final optical density, followed by increased lysis relative to WT cultures (Fig. 2a), like in the absence of cumate (Fig. S7). CRISPRi-mediated silencing of *mreCD* in the WT background (WT + g^mreCD^) slightly reduced growth, closely resembling the growth profile of the Δ*mreB* strain (Fig. 2a), while silencing of *mreBCD* (WT + g^mreBCD^) rapidly and strongly reduced the growth rate and the final optical density of the cultures. When *mreCD* was silenced in the Δ*mreB* background (Δ*mreB* + g^mreCD^), growth reduction occurred more gradually compared to *mreBCD* silencing, but cultures still reached a lower final optical density before lysing. These results demonstrate that loss of the entire *mre* operon (*mreBCD*) has the most severe effect on bacterial growth among all conditions tested.

**Figure 2:**
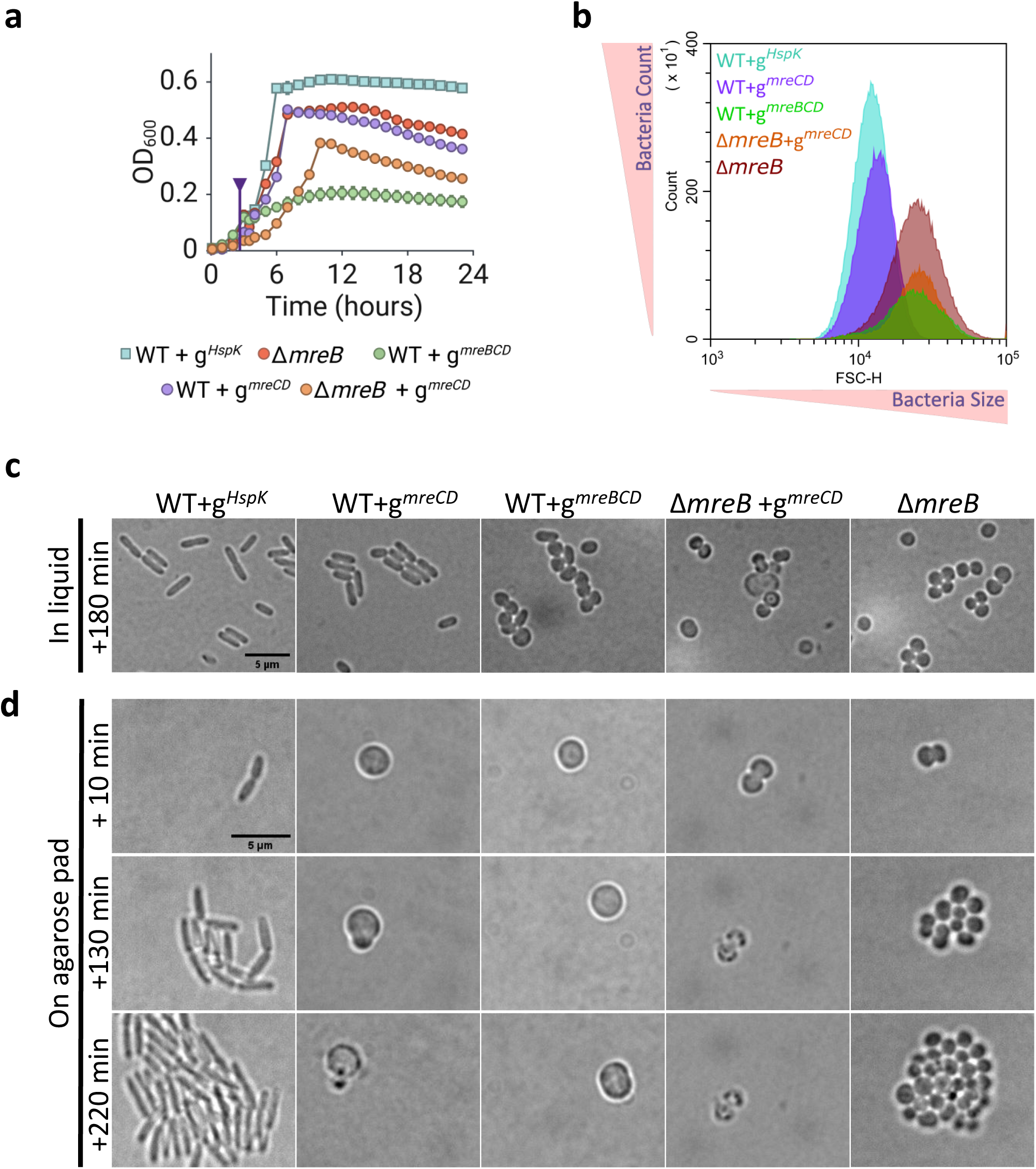
CRISPRi silencing of *mreCD* and *mreBCD* in *P. aeruginosa*. Growth comparison of the Δ*mreB* mutant (brown) and CRISPRi-silenced strains targeting *mreBC* and *mreBCD*. For CRISPRi silencing, cells carried the pFL-dCas9 plasmid along with the pFL-gRNA plasmid expressing guide RNAs (gRNAs) targeting the start codons of the respective ORFs to block transcription elongation. The WT control strain was transformed with a non-targeting gRNA against *HspK*, a heat shock protein gene not present in *P. aeruginosa* (WT + g*^HspK^*, blue). Experimental strains included WT + gRNA targeting *mreBCD* (green), WT + gRNA targeting *mreCD* (orange), and Δ*mreB* + gRNA targeting *mreCD* (purple). **(a)** Growth curves of *P. aeruginosa* strains in LB medium at 37°C, monitored via optical density (OD) at 600 nm over 24 hours. 200 µM cumate was added when the OD_600_ reached 0.1 (purple triangle and vertical line). **(b)** Flow Cytometry Analysis of mreBCD, mreCD silencing by CRISPR-i method and **(c)** their control for morphology with BF images. The figure shows forward scatter height (FSC-H) versus cell count for WT + g^HspK^ (blue), *ΔmreB* (brown), WT+g^mreBCD^ (green), WT+ g^mreCD^ (orange), and *ΔmreB* +g^mreCD^ in *P. aeruginosa* strains after 3 hours of incubation with cumate. FSC-H refers to size changes in bacterial cells, while counts show the cell growth of the population. **(d)** Single-cell imaging for full activation of the dCas9 system. The figure displays time-lapse bright-field images tracking the morphological changes and growth dynamics of WT, *ΔmreB*, g^MreCD^, and g^MreBCD^ *P. aeruginosa* cells over 220 minutes. Scale bar: 5 μm.

To assess changes in cell morphology and population dynamics, we first performed flow cytometry analysis after 3, 5, and 9 hours of incubation with cumate (Fig. S8a, b). 3h after activation of dCas9 by addition of cumate, cell counts of the strains (Fig. 2b) correlated with their differential growth (Fig. 2c). Forward scatter height (FSC-H), which correlates with cell size, was higher in the Δ*mreB* than in the WT control strain, consistent with the wider spherical morphology characteristic of Δ*mreB* mutant cells (Fig. 1c). Both WT+g^mreBCD^ and ΔmreB+g^mreCD^ displayed FSC-H distributions similar to Δ*mreB*, suggestive of shape defects similar to the *mreB* deletion. However, silencing of *mreCD* in the WT background led to a smaller shift in FSC-H values, indicating only mild shape changes and resulting in FSC-H values similar to the WT+g^HspK^ control strain (Fig. 2b**, WT + g^mreCD^**). In agreement with this, microscopic examination showed that 3h after cumate addition, *mreCD*-silenced-cells displayed a rod-shaped morphology, though they were shorter than WT cells (Fig. 2c). In contrast, WT+g^mreBCD^ cells were a mixture of short rods, ovoid (ovococcal), and spherical shapes, whereas Δ*mreB* cells and ΔmreB+g^mreCD^ were virtually spherical (Fig. 2c). Taken together, these results suggested that MreCD may not be required for rod shape in *P. aeruginosa* or, alternatively, that silencing of *mreCD* was less efficient than silencing of *mreBCD.* Silencing efficiency in CRISPRi depends on the sgRNA’s target position, especially if it binds near the promoter and on the non-template strand, as well as its sequence and structure, which together affect how strongly transcription is blocked^44^. 5 and 9 hours post-induction of dCas9, WT and Δ*mreB* cells remained rod-shaped and spherical, respectively, though they were smaller, as expected in the stationary phase, where bacteria reduce their size and metabolic activity^45,46^ (Fig. S8a, b). Cell populations with silenced *mreCD* and *mreBCD* in the WT background still contained mostly short rods and spherical cells, respectively; however, both exhibited increased heterogeneity in cell size and shape, and wild-type-like rod-shaped cells were also observed in the microscopy fields (arrowheads in Fig. S8b). We hypothesized that *Pseudomonas aeruginosa* has a lot of efflux pumps that could expel cumate efficiently thus lowering the intracellular concentration over time, which may have diminished silencing efficiency, causing reduced silencing and allowing for some recovery of cell morphology.

To assess the full phenotypic impact of CRISPRi silencing, we implemented an alternative experimental design in which cells at low density (OD_600_ nm 0.0001) were exposed to cumate for 12 hours in liquid medium, with two additional additions of 200 µM cumate. In these conditions, *mreCD-* and mreBCD-silenced cultures did not grow, whereas WT and Δ*mreB* strains did (Fig. S9). Samples from these liquid cultures were transferred onto fresh 1% LB agarose pads containing the same concentration of cumate to observe surviving cells, and single-cell growth was monitored for more than 3 hours. WT and Δ*mreB* cells kept growing and propagating as rods and spheres, respectively (Fig. 2d). The few *mreBCD-* and *mreCD-*silenced survival cells were enlarged spheres that sometimes further enlarged their volume but were unable to propagate and ultimately lysed (Fig. 2d). Other WT+g^mreBCD^ cells in the field were already lysed (Fig. S10). Taken together, our results indicate that while *mreB* is dispensable for growth, *mreCD* is essential for both rod-shaped and viability in *P. aeruginosa*.

### Deletion of *mreB* disrupts division geometry; co-culturing with WT does not affect their division but does affect cell size

We next monitored the growth of WT cells constitutively expressing mScarletI and the *ΔmreB* mutant expressing sfGFP over time using super-resolution Lattice structural illumination microscopy (Lattice-SIM) (Fig. 3a). Expression of fluorescent proteins was aimed at tracking individual populations in co-culture experiments and better visualise division patterns. Δ*mreB* cells aggregate during growth, which causes errors in the segmentation of cells in phase contrast images (Fig 1a). Control experiments showed that the growth rate of strains expressing the fluorescent proteins was significantly reduced (Fig. S15), as expected from the fitness cost of maintaining plasmids^47^, and cell length increased in both strains (Fig. S13a-c, g) while the width of Δ*mreB* did not change, but WT increased (Fig. S13e, f, d, h).

**Figure 3:**
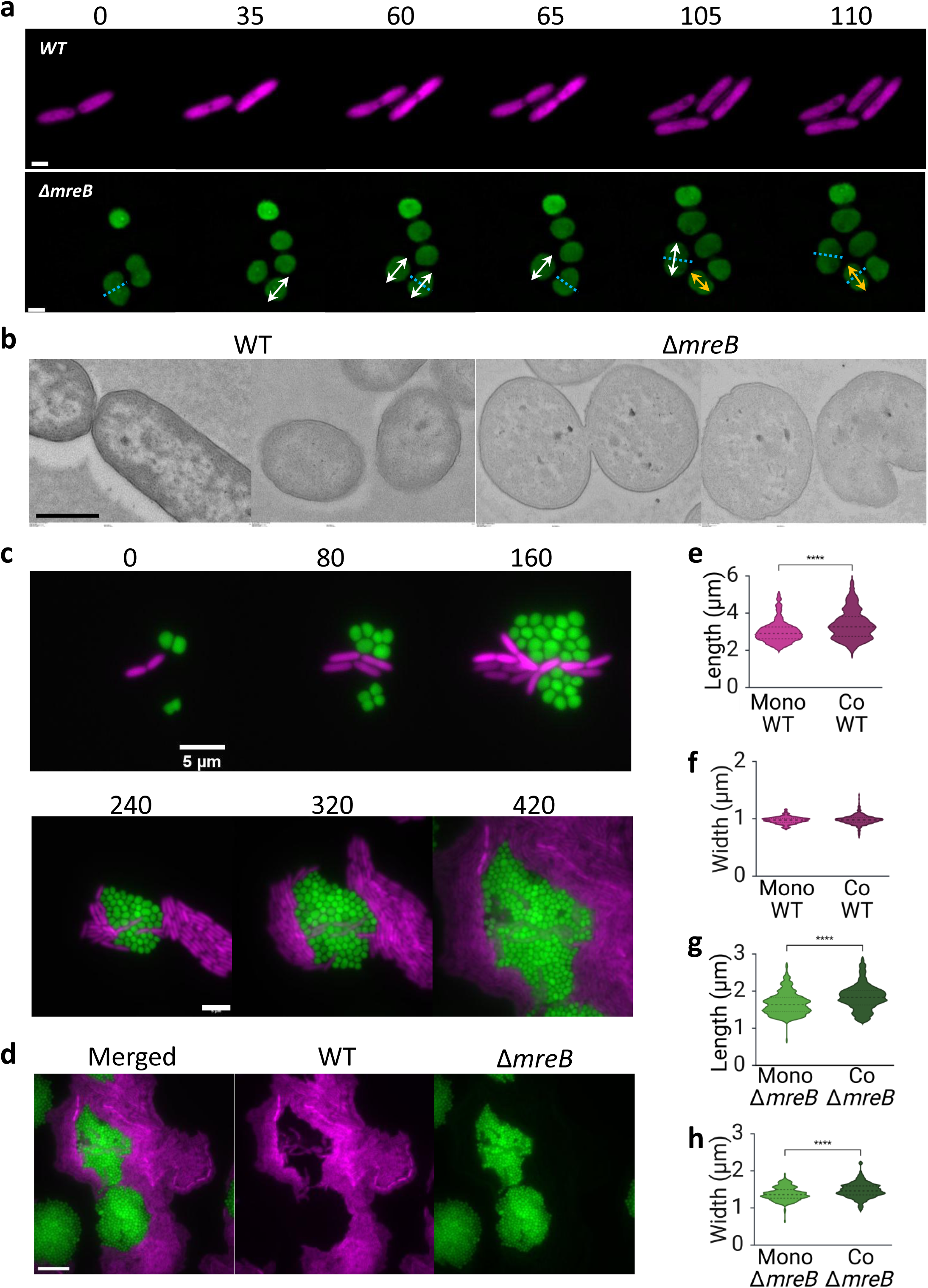
Cell division and growth rate of the *mreB* mutant in mono- and co-cultures. **(a)** Time-lapse Lattice-SIM images of *P. aeruginosa* WT + mScarletI (top row) and Δ*mreB* + sfGFP (bottom row) growing on 1% LB agar pads at 37 °C. Images correspond to 180 minutes (movie acquired at 5-minute intervals for 205 minutes; see **S. Movie 2a, b**). White and yellow arrows indicate the elongation axis of Δ*mreB* cells, while dotted blue lines refer to the division plane on the referred cells. Scale bar: 1 μm. **(b)** Transmission Electron Microscopy (TEM) images of WT and Δ*mreB* cells. Scale bar: 500 nm. **(c)** Time-lapse epifluorescence images of WT + mScarletI (magenta) and Δ*mreB* + sfGFP (green) co-cultured on 1% LB agar pads at 37 °C. Images correspond to 420 minutes (see S. Movie 3a, b). Scale bar: 5 μm. **(d)** Full field of view at the 420-minute time point from **(c)**, with split channels: merged image, WT + mScarletI (magenta), and Δ*mreB* + sfGFP (green). Scale bar: 200 µm. **(e–h)** Quantitative analysis of cell morphology in mono- and co-cultures. ‘Mono F’ refers to fluorescence strain in monoculture, while Co F refers to fluorescence strain in coculture. **(e)** Length and **(f)** width comparison of WT + mScarletI in mono- and co-culture; **(g)** length and **(h)** width comparison of Δ*mreB* + sfGFP. Statistical analysis was performed using the Mann–Whitney U test (unpaired, nonparametric, two-tailed) for all comparisons.

We next performed time-lapse experiments. In rod-shaped cells, division typically occurs at midcell, in a plane perpendicular to the longitudinal axis -and thus to the cell surface-facilitating two-dimensional colony expansion during growth (Fig. 1a, 3a, **WT**). Spherical Δ*mreB* cells exhibited misoriented elongation axes after division and slower growth rates compared to WT (Fig. 1a, 3a**, Δ*mreB***). Δ*mreB* mutants of *P. fluorescens* were recently reported to slightly elongate before dividing in subsequent orthogonal planes^39^, as previously described for the model coccoid bacterium *Staphylococcus aureus*^48^, which contains no *mreB* gene in the genome^17^. In *P. aeruginosa ΔmreB cells,* division planes also shifted by approximately 90° and cells were slightly elongated between two consecutive divisions. In addition to alternating division planes, daughter cells frequently divided asymmetrically over time, resulting in unequal cell volumes in many cases (Fig. S11a), as observed in *P. fluorescens*^39^. Observation of longitudinal and transversal mid-sections of both WT and ΔmreB cells by Transmission Electron Microscopy (TEM) showed that in WT bacteria, septum constriction was symmetrical around the cell circumference (Fig. 3b), as expected from Z-ring constriction at midcell. However, in *mreB* deleted, constriction was often asymmetrical or observed on only one side of the cell, suggesting (Fig. 3b, **Δ*mreB***). The thickness of the envelope of *ΔmreB* cells was similar to that of WT cells (Fig. S11b, c).

In time-lapse co-culture experiments of Δ*mreB* and WT cells (Fig. 3c), Δ*mreB* cells aggregated and did not spread, while WT cells exhibited collective motility and spread around the Δ*mreB* colony. The two cultures did not mix with each other during co-culture, although WT cells were sometimes seen forming connections with Δ*mreB* cells, but actually they were there at the beginning, before cells started growing and spreading on the pad (Fig. 3d). The average length of WT cells increased in co-culture (Fig. 3e, Fig. S14a), but their width remained unchanged (Fig. 3f, Fig. S14b). In contrast, Δ*mreB* cells became slightly longer (Fig. 3g, Fig. S14c) and wider -and thus larger-compared with mono-culture conditions (Fig. 3h, Fig. S14d). The growth rate of *mreB* cells was also slightly increased in co-culture conditions (Fig. S15). These findings suggest that the growth rate and cell dimensions of Δ*mreB* cells are affected in bacterial communities, possibly due to mechanical confinement or environmental cues that *ΔmreB* cells cannot properly process.

### Deletion of *mreB* abolishes motility in *P. aeruginosa*

While WT cells formed well-spread colonies on agar plates, Δ*mreB* cells aggregated into irregular clusters, often growing on top of each other (Fig. 1a, 2d, 3c). The coordinated development of a *P. aeruginosa* colony on an agar plate relies on both cell division and collective bacterial swarming motility, which allows cells to move along the surface^49^. The inability of *ΔmreB* cells to spread efficiently on the agar pad during growth could be due to random cell division orientations relative to the cell surface (Fig 3a, Fig. S11), defects in motility, or both. We next sought to investigate if deletion of *mreB* affected the motility of *P. aeruginosa*.

*P. aeruginosa* employs three distinct modes of motility-swimming, twitching, and swarming-to navigate its environment and establish infections^50^. Swimming motility occurs in liquid environments and is mediated by a polar, helical flagellum, which rotates to propel the cell forward. Twitching motility, observed on solid surfaces (Fig. S12), relies on the extension, attachment to surfaces and retraction of type IV pili^51^, and is crucial for biofilm formation and surface colonization. Swarming motility, occurring on semi-solid surfaces, is a coordinated movement of bacterial populations facilitated by flagella, pili, and the production of tension-active molecules, namely rhamnolipids, which modify the substrate and facilitate the collective moment. This behaviour enables *P. aeruginosa* to form complex branching patterns and is essential for tissue invasion and pathogenicity^53^. It was previously reported that when wild-type *Myxococcus xanthus* cells were exposed to A22, all cells stopped moving within 5 minutes, indicating that a functional MreB cytoskeleton is necessary for *Myxococcus* motility^54^. Also in *P. aeruginosa*, A22-induced inactivation of MreB leads to mislocalization of the type IV pilus retraction ATPase PilT and reduction in motility^22,24,25^. It is important to remember that A22 not only inhibits MreB and alters cell shape, but also affects bacterial signalling via c-di-GMP, disrupts various forms of motility, prevents biofilm formation, and reduces surface adhesion, though it remains unclear which of these effects are directly linked to MreB inhibition^25^. After observing irregular clustering of Δ*mreB* cells at the single-cell level (Fig. 1a and Fig. 3a, c), we first performed twitching, swarming and swimming motility assays on semi-solid agar plates.

In twitching assays (Fig. 4a), the WT strain displayed a distinct halo ranging from 1.5 to 2.0 cm (1001 ±14 mm^2^ total area) after 48 hours of incubation at 37°C. In contrast, no visible twitching halo was observed in the Δ*mreB* mutant (27 ±5 mm^2^ total area), consistent with the proposed role of MreB in type IV pilus function. In swarming assays (Fig. 4b), the WT strain exhibited the characteristic dendritic spreading pattern, indicative of functional, flagella-driven motility^55^. In contrast, Δ*mreB* formed a compact, non-spreading colony, indicating a severe motility defect. In swimming assays, the WT displayed a growth area of 2654 ±28 mm², while Δ*mreB* cells spread only to 396 ±15 mm² (Fig. 4c). To further investigate this defect, we used Scanning Electron Microscopy (SEM) to examine the flagella of *P. aeruginosa* WT and Δ*mreB* cells. In SEM images, WT cells retained their typical rod-shaped morphology and exhibited one or two polar flagella (Fig. 4d, **WT**). While *P. aeruginosa* typically has one flagellum, hyper-swarming variants may exhibit multiple flagella^56^. Similarly, spherical *ΔmreB* mutant cells also displayed one or two flagella (Fig. 4d, **Δ*mreB***), indicating that flagella assembly still occurs in the absence of MreB, though their localization was less predictable due to the loss of defined cell poles in spherical cells. However, the loss of cell polarity in these round cells likely causes a disorganized distribution of flagella on the cell surface, impairing coordinated motility. Taken together, these results demonstrate that both pili and flagella are still present in Δ*mreB* cells, but exhibit reduced or dysfunctional activity, ultimately compromising motility.

**Figure 4:**
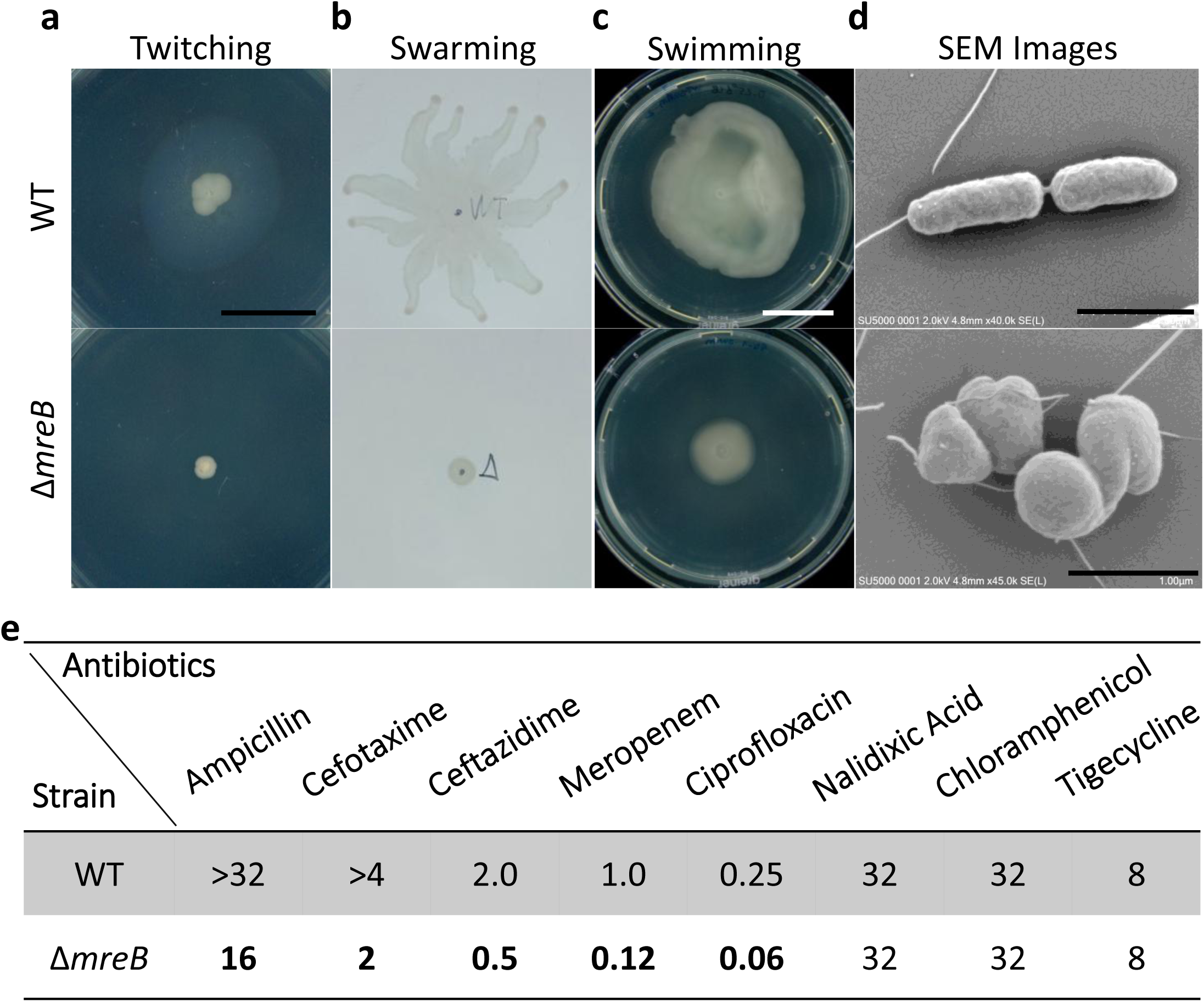
Effect of *mreB* deletion on motility and antibiotic resistance. **(a–c)** Representative images of motility assays for *P. aeruginosa* WT (top) and Δ*mreB* (bottom) strains: **(a)** twitching on solid LB agar, **(b)** swarming on semi-solid LB agar, and **(c)** swimming on semi-solid LB agar. Scale bar: 2 cm. **(d)** Scanning Electron Microscopy (SEM) images of WT (top) and Δ*mreB* (bottom) cells. Scale bar: 1 μm. **(e)** Table of minimum inhibitory concentrations (MICs, mg/L) for a panel of antibiotics tested against WT (PAO1-fix) and Δ*mreB* (PAMNT054) strains. MIC values that decreased in the Δ*mreB* mutant are highlighted in bold.

### Deletion of *mreB* increases the susceptibility of *P. aeruginosa* to antibiotics targeting cell wall synthesis

Next, we tested the sensitivity of the Δ*mreB* mutant to antibiotics targeting several cellular processes. Antibiotics tested included: chloramphenicol and tigecycline (protein synthesis inhibitors), ciprofloxacin and nalidixic acid (DNA replication inhibitors), ampicillin, cefotaxime, ceftazidime and meropenem (PG synthesis inhibitors, β-lactams). The minimum inhibitory concentration (MIC) values for the antibiotics tested in WT and Δ*mreB* strains are listed in Fig. 4e, while their classification and mechanisms of action in *P. aeruginosa* are summarised in Table S2. Chloramphenicol and tigecycline inhibit the 50S and 30S ribosomal subunits, respectively. Chloramphenicol blocks protein synthesis while tigecycline prevents tRNA binding^57^. Sensitivity to either chloramphenicol or tigecycline was not affected in the Δ*mreB* mutant relative to the WT strain (Fig. 4e)

Nalidixic acid and ciprofloxacin belong to the quinolone and fluoroquinolone classes, respectively, and both inhibit bacterial DNA replication by targeting DNA gyrase. Despite their similar mechanism of action, the two antibiotics differ significantly in potency, spectrum of activity, and clinical applications^58^. Typically, ciprofloxacin is more potent and has better penetration than nalidixic acid in *P. aeruginosa* cells. While the deletion of *mreB* did not affect the MIC of nalidixic acid, it rendered *P. aeruginosa* more sensitive to ciprofloxacin (Fig. 4e). The selective sensitivity of Δ*mreB* cells to ciprofloxacin in *P. aeruginosa* may be attributed to several factors. One key reason is the involvement of distinct efflux pump systems: ciprofloxacin is primarily expelled by the MexEF-OprN and MexCD-OprJ systems^59^, while nalidixic acid is largely removed by the MexAB-OprM^60^system. Additionally, their differing chemical charges influence membrane permeability. Ciprofloxacin exists as a zwitterion at physiological pH, while nalidixic acid carries a net negative charge, resulting in differential uptake and retention^61^. Disruption of *mreB* has been shown to alter membrane rigidity and permeability, potentially affecting antibiotic entry and effectiveness^62^.

β-lactams inhibit PG cross-linking by blocking the transpeptidase active sites of aPBPs and bPBPs, thereby inhibiting cell wall synthesis. Δ*mreB* cells exhibited increased sensitivity to ampicillin, cefotaxime, ceftazidime and meropenem, consistent with the role of MreB in PG synthesis (Fig. 4e). Ampicillin is a broad-spectrum antibiotic that primarily inhibits PBP2^63^. Ceftazidime targets preferentially PBP3, with weaker binding to PBP1a and PBP1b, and PBP2^64^. Since Cefotaxime is not used regularly against *P. aeruginosa*, its primary PBP target is unknown, but it is known to have higher affinity for PBP4, PBP1b, and PBP2 in *E. coli,* and for PBP4 and PBP3 in *S. aureus*. Meanwhile, meropenem targets PBP4, PBP3, PBP2, PBP1b and PBP1a, respectively in *P. aeruginosa*^64^ (Table S3). Thus, while these four β-lactams display different affinities towards the different PBPs of *P. aeruginosa,* the two aPBPs, PBP1a and PBP1b, are common targets for all of them^11,64,67^ (Table S3), suggesting an increased requirement of aPBPs activity in Δ*mreB* mutants, as expected in the absence of a functional Rod complex. In rod-shaped bacteria, sidewall synthesis is mediated by the action of the spatially distinct activities of the Rod complex and aPBPs^10^. The Rod complex displays directional motion around the cell circumference and thus inserts PG anisotropically, promoting rod shape, whereas aPBPs diffuse in the membrane and insert PG isotropically^68^. In the absence of *mreB*, and thus a functional Rod complex, aPBPs may be responsible for growth and thus the spherical morphology of *mreB* mutant cells. However, the primary (higher affinity) target of ceftazidime is PBP3, the PBP associated with the divisome, which could also suggest an increased requirement for divisome activity for Δ*mreB* mutants to grow. In agreement with this, *mreB* mutants were originally reported to overproduce PBP3 and PBP1b, the aPBP traditionally associated with cell division, in *E. col*^42^. Furthermore, the lethality of *E. coli ΔmreB* mutants in rich medium can be suppressed by increased levels of FtsZ, which allows mreB mutants to propagate as spheres^34,35^. In *Neisseria elongata*, deletion of the *mreB* operon resulted in spherical cells, albeit accompanied by duplication of a genomic region, possibly leading to overexpression of the division machinery^69,70^. Notably, one of the six *ΔmreB* mutants generated in our study harbours a second, uncharacterized point mutation in *ftsZ* (Table S1). Thus, growth of Δ*mreB* mutant cells may rely on either increased aPBPs activity or increased activity of the septal synthesis machinery, or both.

Quantitative western blot analysis confirmed that FtsZ levels were similar across all our *ΔmreB* mutants compared with the WT strain (Fig. S3b). Thus, propagation of Δ*mreB* mutants in *P. aeruginosa* does not appear to select for progeny that overproduce FtsZ, unlike in *E. coli*^35^. Although similar levels of activity, increased activity of the divisome proteins cannot be excluded in Δ*mreB* mutant cells, we concluded that the growth of *ΔmreB* mutants in *P. aeruginosa* may rely more on compensatory mechanisms involving increased activity of aPBPs

### Deletion of *mreB* promotes surface adhesion and alters biofilm dynamics

*P. aeruginosa* readily transitions from individual cells (planktonic lifestyle) to structured biofilm communities. Biofilm formation enhances *P. aeruginosa* tolerance to antibiotics and host immune defences, making infections particularly difficult to treat and contributing to their persistence in chronic infections.

The compound A22 has been reported to inhibit biofilm formation and reduce the motility in some clinical isolates *P. aeruginosa*, although its anti-biofilm activity is not consistent across all strains^25^. Swimming and swarming motility have been linked to biofilm formation^49^. In *P. aeruginosa, s*warming requires type IV pili, which are also essential for twitching motility, a key driver in the development of mushroom-like structures and microcolonies in mature *P. aeruginosa* biofilms^71^. Our Δ*mreB* strain also exhibited impaired motility (Fig. 4a-d). To investigate the impact of *mreB* deletion on biofilm behaviour, submerged mono-culture biofilms of the WT and Δ*mreB* strains, expressing the fluorescent proteins mScarlettI (red) and sfGFP (green) respectively, were grown in LB medium and monitored over a 15-hour period (Fig. 5a). Although both cultures were inoculated at identical initial densities, quantification of surface-adhered biovolume after 2 hours revealed that the Δ*mreB* strain exhibited approximately threefold higher surface adherence than the WT (Fig. 5b). Biofilm formation assays revealed marked architectural differences between the WT and Δ*mreB* strains. While both strains initially formed rough, irregular biofilms, the Δ*mreB* mutant formed aggregated colonies early, which then progressively gave rise to denser, more compact submerged biofilms compared to the WT. We then quantified total biofilm volume and roughness, a parameter reflecting surface irregularities and structural heterogeneity in biofilm thickness and architecture^72^. No statistically significant differences were observed in total biofilm volume (Fig. S16a) or roughness (Fig. 5c) between WT and Δ*mreB* mono-cultured biofilms until the 15-hour time point. At t=15h, *ΔmreB* biofilm exhibited a significant reduction in roughness, indicating the formation of a more compact and homogeneous surface compared to WT biofilms. To assess biofilm temporal dynamics, we then calculated the biofilm growth rate (µm³/min) at each time point over the 15-hour period. Both strains exhibited positive growth rates during the initial phase. WT biofilms reached maximum growth rate at around 6 hours, whereas Δ*mreB* biofilms peaked slightly earlier, at about 4.5h, followed by a gradual decline (Fig. 5d). Notably, after 6 hours, WT biofilms showed a sharp decline and reached negative growth rate values at around 7.5 hours, suggesting biofilm dispersal, detachment or degradation at this stage. In contrast, Δ*mreB* biofilms progressively decreased their growth rate for about 10.5 hours and then stopped growing, but did not display negative growth rate values suggestive of biofilm dispersal (Fig. 5d). Taken together, our results show that the deletion of *mreB* enhances early surface colonization and inhibits biofilm dispersal, promoting the formation of more stable and persistent biofilm compared to the WT. These findings have important implications for antibiotic susceptibility and host-pathogen interactions, as biofilm structure and density are closely linked to bacterial persistence, immune evasion, and the potential for chronic infection.

**Figure 5:**
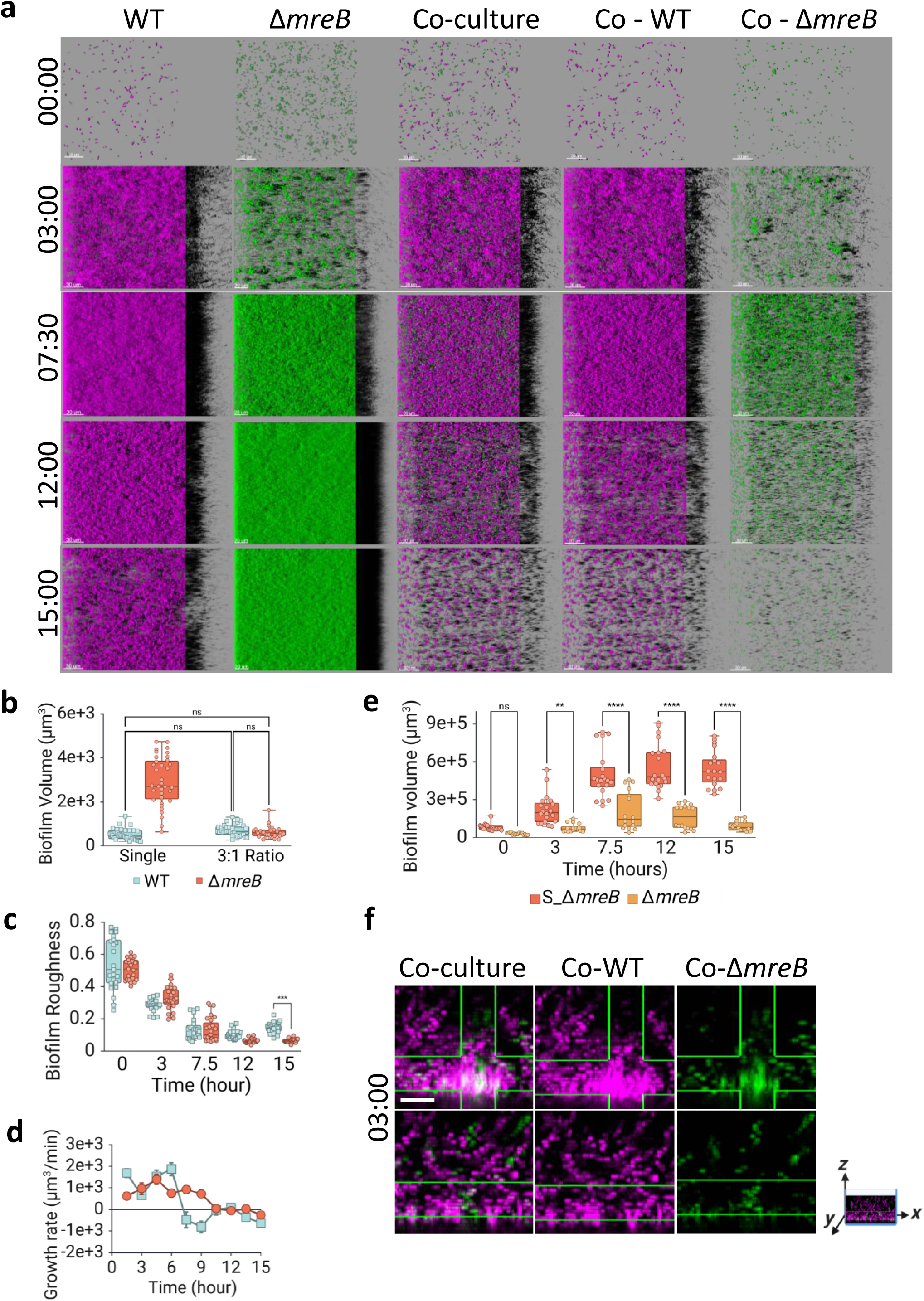
Growth dynamics and structure of wild-type (WT) and *ΔmreB* biofilms. **(a)** Time-lapse confocal microscopy images showing biofilm development of *P. aeruginosa* WT + mScarletI (magenta) and Δ*mreB* + sfGFP (green) strains under monoculture and co-culture conditions over 15 hours. Scale bar: 30 μm. **(b)** Quantification of biofilm volume at the adhesion stage (t = 0) for WT + mScarletI and Δ*mreB* + sfGFP strains in monoculture and in co-culture (3:1 ratio, WT:Δ*mreB*). Δ*mreB* exhibits significantly greater initial biofilm volume than WT in monoculture, whereas in co-culture, both strains begin with similar adhesion volumes. **(c)** Biofilm surface roughness and **(d)** biofilm growth rate of WT (blue) and Δ*mreB* (orange) monocultures over 15 hours. **(e)** Comparison of Δ*mreB* biofilm growth in monoculture versus co-culture (3:1 WT:Δ*mreB*), showing reduced growth in the presence of WT cells. Scatter points represent individual biological replicates. **(f)** Confocal z–y section of a co-culture biofilm at the 3-hour time point, showing merged and split fluorescence channels of WT + mScarletI and Δ*mreB* + sfGFP strains. Scale bar: 25 μm. Statistical analysis: Two-way ANOVA with Tukey’s multiple comparisons test was used for biofilm volume at adhesion **(b)**, and Two-way ANOVA with Bonferroni correction was applied for surface roughness **(c)** and Δ*mreB* biofilm growth comparison in mono- and co-culture **(e)**.

### Δ*mreB* mutant exhibits impaired biofilm growth when co-cultured with wild-type cells

In natural and clinical environments, *P. aeruginosa* often exists in complex biofilms where diverse microbial populations compete for space, nutrients and survival^73^. Within these communities, competition is shaped by differences in surface colonisation efficiency, motility, and growth rates^2,56^. Using a combination of computational simulations and experimental validation, it has been proposed that individual cell shape drives spatial organisation within biofilms^26^. A reported model predicts that in a co-culture biofilm, different cell shapes self-organize into layered structures, and that spherical cells rise to the biofilm surface while rod-shaped cells are better at colonizing the basal surface and edges, irrespective of nutrient availability^26^. To assess how loss of *mreB* affects spatial organisation and fitness within a genetically heterogeneous biofilm community, we performed competition experiments by co-culturing differentially labelled WT and *ΔmreB* strains in the same biofilm. Given the previously observed threefold difference in adhesion levels (Fig. 5b), we first determined the appropriate WT:Δ*mreB* inoculum ratio to normalize initial surface colonization (Fig. S16b). A 3:1 WT to Δ*mreB* ratio was selected, resulting in comparable surface-adhered biovolume of bacteria at t0 (Fig. 5a, **Co-culture**). Under these conditions, WT biofilm growth remained unaffected by the presence of the Δ*mreB* strain relative to mono-culture conditions (Fig. 5a and Fig. S16c).

In contrast, Δ*mreB* biofilm growth was significantly impaired in co-culture, exhibiting a sharp reduction in volume and growth rate relative to Δ*mreB* mono-culture (Fig. 5a-e). Confocal side projections (Fig. 5f) showed that WT and Δ*mreB* cells were intermixed throughout the biofilm, contradicting the model prediction of layer segregation^26^. Rod-shaped cells (WT) did not displace spherical (Δ*mreB)* cells from the base of the biofilm. Instead, aggregated Δ*mreB* cells were frequently observed at the base of the biofilm, overlaid by WT cells (Fig. 5f), a spatial arrangement also seen in long-term single-cell time-lapse experiments (Fig. 3c,d). This pattern occurred regardless of nutrient flow direction, suggesting that the positioning of *ΔmreB* cells at the biofilm base was not driven by nutrient gradients but rather by their physical association with the surface. Although the presence of spherical mutant cells in the upper layers of biofilms has been reported in the literature, the motility ability of these mutants remains poorly characterized^26^. It has been reported that GFP fluorescence can be reduced in biofilms due to limited oxygen availability, potentially affecting fluorescence detection in deeper layers^74^. To exclude the possibility that differences in fluorescent proteins or plasmid backbones affected biofilm formation, we swapped the plasmids between the strains, introducing mScarletI into the Δ*mreB* mutant and sfGFP into the WT strain, and monitored their growth again. No noticeable effect of the fluorescent proteins was observed during either the adhesion step (Fig. S16e) or during 15 hours of biofilm growth (Fig. S16d). Based on our observations, the restricted vertical movement of *ΔmreB* cells is more likely due to their limited motility than to nutrient accessibility. Taken together, our findings demonstrate that deletion of *mreB* not only affects biofilm structure in monoculture but also compromises the competitive fitness and spatial dynamics of *P. aeruginosa* in mixed biofilm communities.

### *P. aeruginosa* natural isolates can carry deletions of *mreB*

With the exception of the closely related *P. fluorescens*^39^, deletion of *mreB* has been reported to be lethal in other rod-shaped bacterial species where this has been investigated. The viability and fitness of the Δ*mreB P. aeruginosa* mutant generated in this study under standard laboratory conditions raised the intriguing question of whether loss of *mreB* might also occur naturally in *P. aeruginosa* populations in the wild. Phylogenetic studies indicate that cocci evolved from rods and thus that coccoid morphology is an evolutionary loss-of-function trait^75,76^. In support of this, a rod-to-coccus transition was reported to be selected during the evolutionary history of the pathogenic *Neisseria and Moraxellaceae* families, likely as an adaptation to colonization of the nasopharynx – portal for infections and ecological niche of many pathogens^69^. Notably, genetic changes associated with this morphological change in the *Neisseria* family included deletions encompassing the elongation machinery, including the *mreBCD* operon^69^. To explore whether natural *P. aeruginosa* isolates carry deletions of *mreB* or mutations in the sequence of *mreB*, we screened the core genome sequences of 10167 strains from the NCBI collection by using the *mreB* sequence from strain PA96 (GenBank: CP007224.1) as a reference. Remarkably, we identified 18 strains (0.18%) harboring *mreB* alleles encoding truncated MreB proteins at various positions. Protein sequences of these strains were listed in (Table S4). The discovery that 0.18% of natural *P. aeruginosa* isolates harbour truncated *mreB* alleles demonstrates that the loss/changes of this traditionally “essential” gene can occur and persist in natural populations.

To determine whether alterations in the *mreB* sequence affect cell morphology, we examined the cell shape of four^77^ representative clinical isolates among the 18 strains identified (Fig. 6). Quantitative morphometric analysis (Fig. S18a-d) revealed that four of these strains exhibited significantly reduced cell length compared to wild-type *P. aeruginosa*, whereas one strain displayed increased length. Notably, all strains with reduced length showed circularity values approaching 1, in contrast to the wild type, which retained a more elongated rod-like morphology. Importantly, strains carrying truncated *mreB* exhibit morphological behaviours resembling those of Δ*mreB* mutants, indicating that these truncations functionally phenocopy *mreB* deletion at the level of cell shape. Overall, our results indicate that natural variation in *mreB* produces distinct cell shapes in *P. aeruginosa* and that partial loss of elongation function can be maintained in natural populations.

**Fig. 6:**
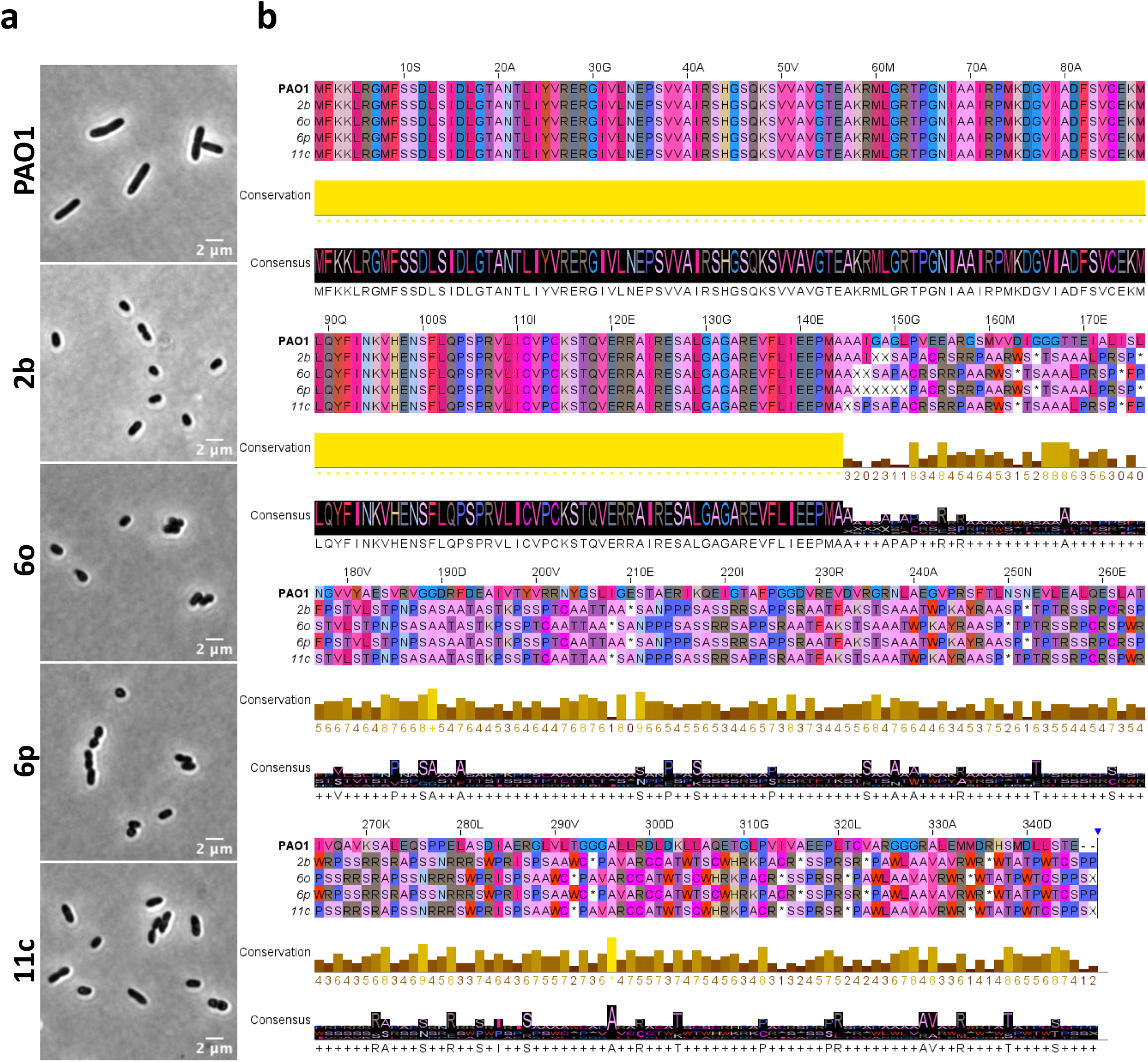
Phase-contrast images of *P. aeruginosa* PAO1 (WT), four clinical isolates carrying truncated mreB and their amino acid sequence comparison. **(a)** *P. aeruginosa* PAO1 (wild type) and four clinical strains carrying truncated **mreB** (2b, 6o, 6p, 11c) were grown in LB rich medium until an OD₆₀₀ of 0.5 and were imaged. All clinical isolates are shorter than WT. **(b)** Alignment of clinical isolates with reference sequence of PAO1 shows 100% conservation until 145^th^ amino acid (aa). After that point aa sequence of clinical isolates are similar to each others while different from reference sequence. Colour selection for aa sequence is gecko Sunset. Besides protein sequence, their conservation and consensus scores are listed below the sequences to which they belong.

## Discussion and Conclusion

Our findings demonstrate that deletion of *mreB* in *P. aeruginosa* profoundly alters several aspects of bacterial physiology, including cell morphology, motility, antibiotic susceptibility, and biofilm formation. Unlike in *E. coli*, where *mreB* is essential unless suppressor mutations arise or FtsZ is overproduced, *P. aeruginosa ΔmreB* mutants remain viable without apparent compensatory mutations or upregulation of FtsZ, indicating a different physiological tolerance for loss of the elongasome. This viability may be supported by increased activity of PG-synthesizing enzymes outside of the MreB-guided elongation system, such as bifunctional aPBPs and divisome-associated enzymes, particularly PBP3. Consistent with this, Δ*mreB* cells exhibited increased sensitivity to β-lactam antibiotics, which target PBPs. This hypersensitivity implies that growth in the absence of MreB depends on elevated PBP activity, likely via PBP1a or divisome-associated enzymes.

*ΔmreB* mutant cells exhibit a distinct spherical morphology, which is a hallmark of the loss of MreB’s involvement in maintaining rod shape. Our data further reveal that *mreB* deletion exerts a strong polar effect on downstream genes *mreC*D, significantly reducing MreC protein levels-an effect not previously demonstrated in *P. aeruginosa*. While MreB is dispensable for survival, our CRISPRi experiments confirm that *<u>mreCD</u>* <u>are essential</u>: strong knockdown led to cell rounding, growth arrest, and cell death. These results suggest that the MreBCD complex, and particularly MreC and MreD, are crucial for both the maintenance of rod shape and the survival of *P. aeruginosa*, and that the deletion of mreB does not fully compensate for the essential functions of the *mreCD* operon. Therefore, while mreB is dispensable for growth under certain conditions, the mreCD operon plays an irreplaceable role in ensuring the structural integrity and survival of this bacterium. Importantly, our study does not allow to conclusively demonstrate whether the spherical phenotype of the Δ*mreB* mutant is due solely to the absence of MreB or to decreased MreCD expression. However, the fact that *P. aeruginosa* cells treated with A22, which disrupts MreB polymerisation, also become spherical supports the conclusion that MreB itself is critical for maintaining rod shape^22,24,25^. Further investigations are needed to determine whether the polar effect of Δ*mreB* on MreC is due to direct transcriptional regulation or a secondary consequence of *mreB* loss. And still, the specific role of MreC and MreD in cell morphology remains unclear, as the effects of *mreD* silencing or deletion alone have not yet been determined.

In addition to increased cell width and spherical shape, Δ*mreB* cells exhibited a reduced growth rate compared to wild-type cells, suggesting the functional significance of MreB in maintaining both shape and optimal cellular dynamics during growth. The findings also highlight differences in cell volume, with *ΔmreB* cells exhibiting larger, more heterogeneous volumes.

In rod-shaped *P. aeruginosa*, the spatial organization of cell division is highly regulated, occurring perpendicularly to the long axis of the cell to allow for predictable and symmetric binary fission. In contrast to rods, spherical bacteria can, in principle, divide in any midcell plane and still generate identical daughter cells. Our Lattice-SIM imaging of Δ*mreB* mutants revealed that consecutive divisions frequently occur in orthogonal planes. Interestingly, similar division patterns have been reported in coccoid bacteria such as *S. aureus*^48^, which naturally lack *mreB*, and in Δ*mreB Pseudomonas fluorescens* cells, which exhibit rotational division planes with occasional asymmetry^39^. *E. coli* cells treated with mecillinam or deleted of rodA, which induce cell rounding, also formed division rings in random perpendicular planes between the nucleoids^78^. Divisions in different planes may contribute to geometric disorganization and cell aggregation within microcolonies. Interestingly, our TEM images support the idea that, in Δ*mreB*, constriction at the division site does not begin equally on both sides of the cell, leading to asymmetrical division. This asymmetrical constriction pattern is reminiscent of previous observations in *B. subtilis*, *E. coli* and *Cyanophora paradoxa,* where insufficient FtsZ availability or improper Z-ring formation results in partial arcs rather than complete rings, leading to asymmetric initiation of cell division^79^.

Beyond its role in morphogenesis, we found that MreB is essential for motility in *P. aeruginosa*. Previously, in *Myxococcus xanthus*, MreB has been implicated in controlling motility by coordinating cytoskeletal elements with the pilus machinery^54^. Also, prior work showed that A22 treatment disrupts swarming motility modes in *P. aeruginosa* by altering pilus localisation^24^. Our findings in *P. aeruginosa* are consistent with the production of non - functional pili. Furthermore, we observed a decrease in swimming motility in *ΔmreB* cells, indicating non-functional, mislocalised, or lost flagella. TEM and SEM images reveal that WT cells have flagella, which are specifically positioned at the cell poles, ensuring proper directional movement, while Δ*mreB* cells have flagella but are mostly mislocalised and non-functional. This suggests that MreB is not required for pili and flagella biogenesis per se but is likely essential for proper positioning or function of the motility apparatus. By determining rod shape, MreB plays an active role in establishing and maintaining polar identity, linking morphogenetic processes with polar-related cellular processes.

The disorganised division and lack of motility in Δ*mreB* had functional consequences for colony architecture. Δ*mreB* cells were often observed to form dense, amorphous clusters with little outward spreading. In time-lapse co-culture experiments, WT cells continued to proliferate and spread, whereas Δ*mreB* cells remained largely immobile, forming compact aggregates surrounded by WT cells. These observations suggest that the impaired spatial division of Δ*mreB* mutants, the non-functional flagella, and the altered pilus contribute directly to their reduced ability to spread effectively on semi-solid surfaces.

Interestingly, co-culture did influence the morphology of Δ*mreB* cells. While WT cells maintained their width dimension while becoming longer, Δ*mreB* cells became significantly larger, both in length and width, when grown alongside WT counterparts. This suggests that while Δ*mreB* cells do not regain directional growth, their size may be modulated by physical confinement or mechanical interactions with rod-shaped neighbours. But this observation argues against simple mechanical compression as the primary mechanism affecting cell size. The behavior of *ΔmreB* cells in co-culture shows some interesting patterns. During co-culture biofilm formation, *ΔmreB* mutants exhibit significantly impaired biofilm growth compared to their monoculture performance. They tend to remain at the base of the biofilm, overlaid by wild-type cells, suggesting they may indeed be experiencing stress from the surrounding community. The division patterns of *ΔmreB* cells are already abnormal even without WT cells present - they show asymmetrical division and irregular division planes. When examining *ΔmreB* cells through TEM, the constriction at division sites is often asymmetrical and doesn’t begin equally from both sides of the cell. This inherent division defect could be exacerbated by the presence of WT cells. The slightly improved growth rate of *ΔmreB* cells in co-culture compared to monoculture suggests that while these cells may be under stress, they are not completely inhibited from dividing. However, their continued enlargement could indicate problems with proper timing or completion of cell division, possibly due to the combination of their inherent division defects and the physical constraints imposed by surrounding WT cells.

Therefore, rather than simple mechanical pressure making cells shorter, the data suggest a more complex scenario in which *ΔmreB* cells face challenges completing proper cell division while simultaneously responding to environmental cues from neighbouring WT cells. The enlarged cell size could represent a stress response where cells continue to grow but fail to complete division efficiently, rather than direct mechanical compression

The morphology and motility defects of *ΔmreB* mutants also had clear consequences for biofilm behaviour. In monocultures, Δ*mreB* cells formed dense, highly adherent biofilms that adhered to surfaces nearly three times more efficiently than WT cells. This hyper-adherence may reflect a lack of motility-driven dispersion, leaving Δ*mreB* cells fixed at the surface. However, in mixed biofilms with wild-type cells, Δ*mreB* strains were markedly outcompeted. This suggests that motility is not only dispensable for initial surface attachment but also critical for niche exploration, spatial structuring, and competitive fitness within biofilm communities. Notably, we did not observe the vertical layering of spherical versus rod-shaped cells predicted by recent computational models^26^. This discrepancy may arise from model assumptions that excluded key biological factors, such as the motility comparison of spherical mutant and WT. Besides, their mutant had the same growth rate as WT, which is not the case in *mreB* deletion, even in co-culture, the growth rate was slightly increased, but not close to WT at all. Our results suggest that strong surface attachment of spherical *ΔmreB* cells may inhibit upward movement and layering within the biofilm, altering the predicted spatial dynamics. To fully understand its role in multispecies communities, future studies are needed to assess how *ΔmreB* behaves in co-culture biofilms with other bacterial species.

Given the essential role of the MreBCD complex in maintaining cell morphology and coordinating PG synthesis, it has emerged as an attractive target for novel antibacterial strategies against *P. aeruginosa*. Recent research has identified several compounds that specifically disrupt MreB function, including a novel indole derivative that impairs bacterial growth by targeting MreB polymerization. However, its efficacy is limited by active efflux via the MexAB-OprM system^81^, highlighting the importance of overcoming intrinsic resistance mechanisms. Other studies have identified MreB-derived antimicrobial peptides that directly interact with bacterial membranes, offering an alternative mode of action with promising specificity. TXH11106, a third-generation MreB inhibitor, demonstrates enhanced activity across multiple Gram-negative pathogens, suggesting the potential for broad-spectrum application^82^. Beyond MreB, high-throughput screening approaches have begun to spotlight MreC and MreD as critical nodes in biofilm-specific vulnerabilities, revealing new paths for anti-biofilm therapy. Additionally, the interplay between MreB and type IV pili, regulated by MreB-dependent positioning, underscores a multifaceted role for MreB in virulence and persistence. These findings position the MreBCD complex not only as a key regulator of essential cellular processes but also as a focal point in developing next-generation antimicrobials aimed at combating *P. aeruginosa* formidable defence systems.

In conclusion, our study provides a comprehensive analysis of MreB function in *P. aeruginosa*, demonstrating its pivotal roles in regulating cell morphology, motility, antibiotic response, and biofilm dynamics. Future studies should focus on dissecting the molecular mechanisms of MreB regulation, including the basis for the polar effects on *mreC* and *mreD*, and the basis for MreB-mediated inhibition of cell motility. In addition, exploring MreBCD-associated pathways as antimicrobial targets could uncover novel strategies for combating multidrug-resistant *P. aeruginosa* infections, since some strains in nature harbour mutant MreB proteins.

## Materials and Methods

### General methods and bacterial growth conditions

The *P. aeruginosa* and *E. coli* strains used in this study are listed in **Table S5**. Plasmids and primers used in this study are listed in **Table S6** and **Table S7,** respectively. Strains were grown at 37°C in lysogeny broth (LB) medium, supplemented when required with 50 mg/L Gentamicin (Gm), 100 mg/L Tetracycline (Tet), 25 mg/L Trimethoprim (Tim), 200 µM cumate (Cum), 2 mM theophylline (Teo), and 200 mg/L DAP. For liquid culture growth monitoring, overnight cultures of WT, Δ*mreB*, and *mreBCD*/*mreCD*-silenced strains were diluted to OD₆₀₀ 0.005 in LB medium and inoculated in 96-well plates in triplicate. OD₆₀₀ measurements were recorded every 5 minutes for 24 hours at 37°C, with agitation, using a BioTek plate reader (Agilent, USA). In CIRSPRi experiments, when cultures reached OD₆₀₀ 0.1–0.2, 200 µM cumate was added to one triplicate to activate the dCas9 system, while the second triplicate received an equal volume of LB + ethanol mixture (cumate solvent) as a control.

### Competent *E. coli* DH5α, RHO3 and *P. aeruginosa PAO1-fix* and *ΔmreB* cell preparation

overnight (ON) cultures were first grown in LB medium, with diaminopimelic acid (DAP; 200 µg/mL) added for RHO3. Prior to starting the preparation, 150 mL each of ice-cold 100 mM CaCl₂ and 100 mM CaCl₂ supplemented with 15% glycerol were prepared and stored on ice. The centrifuge was pre-cooled to 4°C. For transformation-competent cell preparation, 1 mL of ON culture was inoculated into 100 mL of LB (plus 200 µg/mL DAP for RHO3) and incubated at 37°C with shaking at 170–250 rpm until reaching OD₆₀₀ nm 0.4–0.5. The cultures were then transferred into two 50 mL Falcon tubes and placed on ice, after that all steps were performed under cold conditions. Cells were pelleted by centrifugation at 4,500 rpm for 5 minutes at 4°C, and the supernatant was discarded. Pellets were gently resuspended in ice-cold 100 mM CaCl₂, with frequent cooling on ice to prevent warming. The resuspended cells were incubated on ice for at least one hour. Following a second centrifugation step under the same conditions, the supernatant was removed, and the cells were resuspended in ice-cold 100 mM CaCl₂ containing 15% glycerol. After another 1-hour incubation on ice, pre-labelled sterile 1.5 mL microcentrifuge tubes were chilled at −20°C. Aliquots of 500 µL were dispensed into each tube and immediately flash-frozen in liquid nitrogen for 5 minutes. The competent cells were then stored at −70°C until use.

### Plasmid uptake by the electroporation method

For electroporation, chemically competent cells stored at −70°C were thawed on ice. A total of 100 µL of competent cells was gently mixed with 5–7 µL of plasmid DNA in a pre-chilled 0.2 cm electroporation cuvette. Electroporation was performed using an Eporator (Eppendorf) at 2.5 kV/cm with a 5-millisecond pulse duration. Immediately after the pulse, 900 µL of LB medium was added directly to the cuvette to facilitate recovery, and the mixture was pipetted several times to ensure complete resuspension before transferring it to a sterile 2 mL microcentrifuge tube. Cells were incubated for 2 hours at 37°C with shaking at 170 rpm to allow phenotypic expression of the antibiotic resistance marker. Following recovery, 50–100 µL of the culture was plated onto LB agar containing the appropriate antibiotic for selection. The remaining volume was centrifuged, resuspended in 100 µL of LB, and also plated onto LB-antibiotic plates to maximize transformant yield. Plates were incubated at 37°C for 24–48 hours. Two to three resulting colonies were selected and assessed for strain morphology to confirm successful transformation and viability.

### Strain constructions

Strain PAMNT054, *mreB* deletion from chromosomal locus tag was generated as follows. The *mreB* deleted 500 bp sequence was designed, and a fragment (Frt_pEX19Gm_for_mreB_deletion) was created by GeneCust. This fragment (**Table S7**) was inserted into the pEX19Gm plasmid^83^ by ligation. The fragment inserted plasmid was transferred to the competent DH5α and then the RHO3 conjugation strain. PAO1-fix grown on SOB agar at 42°C and pEX19Gm included RHO3 strains were conjugated on a SOB plate (supplemented with 200 mg/L DAP), and plates were incubated O/N at 37°C. The next day, grown colonies were transferred to LB plates containing 100 mg/L Gm and 25 mg/L Tim, and incubated at 37°C for O/N. Grown colonies were transferred to LB +25 Tim liquid cultures to permit a second recombination and resolve the merodiploid. Liquid cultures were inoculated at 37°C for O/N. From the O/N cultures, serial dilutions (x10, x100, x1000) were carried out, and 100 µL of each dilution was spread onto LB agar containing 5% sucrose + 25 mg/L Tim and incubated O/N at 37°C. The next day, grown colonies are duplicated onto LB agar plates supplemented with 25 mg/L Tim and 25 mg/L Tim + 100 mg/L Gm and incubated at 37°C for O/N. Double recombinants, whether they were wild-type or the desired mutants, will have lost the vector backbone and would not grow on the plates containing gentamicin. If the colony did not grow on the LB 25 mg/L Tim + 100 mg/L Gm plate, the ON liquid LB culture from the LB + 25 mg/L Tim plate was prepared. DNA was extracted from the selected strains, and *mreB* deletion was tested by PCR using primers *mreBR* and *mreBF* (Table S7). 10 out of 11 clones tested did not contain the *mreB* gene in their genomes.

For fluorescent labelling, plasmids pSEVA627M mScarletI and pSEVA627m sfGFP were transformed into the wild-type *PAO1-fix* and the *ΔmreB* (PAMNT054) strains, respectively, generating strains WT+ mScarletI and *ΔmreB* + sfGFP via 2.5 V electroporation. Electroporation was done on competent cells of *PAO1-fix* and *ΔmreB* strains.

### CRISPR-i system

The expression of the dCas9 gene was achieved under dual transcriptional-translational control, regulated by cumate and theophylline (Fig. S5a). In our system, dcas9 expression was under positive cumate-responsive transcriptional regulation and negative theophylline riboswitch-mediated translational control (Fig. S6 and Fig. S5b). The regulatory module consists of a constitutively expressed cymR gene, encoding the transcriptional regulator CymR, under the control of the P43 promoter (BBa_K143013) and adjacent to the CuO operator site. The CuO operator site is located downstream of the strong Pveg promoter (BBa_K316001), which itself controls the synthesis of a theophylline aptamer (BBa_K3015007)^84,85^. DNA fragments were either synthesized or PCR-amplified from the cumate-inducible expression vector pCT5-bac2.0^86^ and inserted upstream of the dCas9 gene. All constructs were initially made in *E. coli* before being transferred into *P. aeruginosa* PAO1-Fix. The cymR_CuO-RSWtheo regulatory module was substituted for the original aTet-inducible expression system present in the *P. fluorescens* replicative plasmid pPFL-dcas9^21^. The resulting plasmid pPFL-dcas9v2 (in this study) was transformed into *PAO1-Fix* by electroporation and maintained in the presence of tetracycline (100 µg/ml).

The constitutive expression of the guide RNAs was carried out from plasmid pPFL-gTarget, which is selectable in the presence of gentamicin (50 µg/ml)^21^. The targeting of genes was achieved by designing 20-base-pair sequences complementary to the target DNA and located in proximity to the Cas9 protospacer-adjacent motif (PAM) NGG (Table S7). The 20 bp target sequences were PCR-amplified into the pPFL-gTarget vector by bridging a 70 bp oligo comprising a 20 bp guide sequence, flanked by 25 bp of sequence homologies (Table S7, gmreB1 and gmreC1), with the pPFL-gTarget vector at each site of insertion. To do so, the pPFL-gTarget vector was first PCR-linearized at the insertion site using primers PO-106 and PO135 (Table S7) and a high-fidelity polymerase (Q5 from NEB). The amplified 4 kb DNA fragment was then gel-purified and used to insert the 20 bp target sequence using the single-stranded bridging methodology developed by NEB (NEB HiFi DNA assembly). Constructs were first transformed in a chemically competent *E. coli* DH5α strain, selecting for resistance to gentamicin 10µg/ml. Individual clones were selected for plasmid DNA extraction, followed by DNA sequencing to assess the presence of the guide sequence. Positive constructs were then transformed into electrocompetent PAO1-fix strain carrying pPFL-CuO-dcas9, selecting for resistance to tetracycline (100 µg/ml) and gentamicin (50 µg/ml).

### Whole genome sequencing

The same process was applied as previously described^87^. DNA-seq libraries were prepared using Eurofins Illumina’s protocol and sequenced on a NovaSeq 6000 (2 × 150 bp paired-end mode) with ∼5 million reads per sample. Read quality was assessed with FastQC (v0.11.9). Reads were processed in Galaxy (https://galaxy.migale.inrae.fr/) using the default parameters. De novo assembly was performed with Unicycler (v0.4.8.0), and assembly quality was evaluated using Quast (v5.0.2). Variant calling was performed using the bacterial SNP-calling pipeline Snippy (v4.6.0) with default parameters (https://github.com/tseemann/snippy). To identify single-nucleotide polymorphisms (SNPs) and insertions/deletions (indels), the reads from six *mreB* deletion mutants (PAMNT043, PAMNT044, PAMNT048, PAMNT049, PAMNT051, and PAMNT054) were mapped against the wild-type genome PAMNT069 (derived from the PAO1 strain). This analysis was conducted to confirm that no unintended mutations were introduced during the construction of the mutants, aside from the intended *mreB* deletion.

### Genome comparison of *P. aeruginosa strains*

To identify potential truncated MreB proteins occurring in nature, 10167 RefSeq assemblies were retrieved from the NCBI database. The *mreB* gene from the PA96 assembly (CP007224.1) was used as a reference to screen the assemblies using KM^88^, which performs a BLAST-like alignment of assembled genomes. The resulting *mreB* sequences were retrieved from the output, translated into protein sequences, and aligned to detect premature stop codons that would result in truncated MreB proteins and clustered using CD-HIT at 100% identity to remove redundancy. During this step, CD-HIT flagged several sequences as “invalid,” which often corresponded to proteins containing multiple internal stop codons.

### Western Blot

Pre-cultures of *P. aeruginosa* were grown to an OD_600_nm of 1.0, then 1 mL of culture was pelleted and resuspended in 50 µL of resuspension buffer, followed by incubation at room temperature (RT) for 10 minutes. Lysis was induced by adding 50 µL of lysis buffer and incubating on ice for 20 minutes. After lysis, 17 µL of the lysate was transferred to a fresh tube, mixed with 3 µL of Laemmli buffer, and heated at 95°C for 15 minutes. Samples were centrifuged at 10,000 g for 1 minute to remove debris, and the supernatant was collected for analysis. Proteins were separated using 4–15% Mini-Protean TGX precast gels (Bio-Rad) in Tris-Glycine running buffer at 120V for 1 hour, then transferred onto a nitrocellulose membrane using the iBlot 2 system (Invitrogen). The membrane was washed three times in 1X TBST, blocked in 1X TBST + 5% milk for 1 hour, and incubated overnight with the primary antibody (anti-MreC, 1:10,000) at RT under slow agitation. The next day, after three TBST washes, the membrane was incubated with the secondary antibody (anti-rabbit-HRP, 1:10,000) for 2 hours, washed again, and prepared for signal detection.

### Single-cell imaging and analysis

Single-cell imaging was performed using Lattice-SIM with bright-field acquisition. Overnight cultures were diluted to OD₆₀₀ 0.01 in LB medium and grown to OD₆₀₀ 0.15–0.20 (exponential phase, clinical strains with truncated *mreB* alleles were grown until OD600 0.5). Then, 3 µL of culture was placed on a 1% agarose pad and imaged every 5 minutes for 41 cycles at 25% light intensity to observe growth and morphological changes.

For full activation of the dCas9 system, overnight cultures of WT, Δ*mreB*, and *mreBCD/mreCD*-silenced strains were diluted to OD₆₀₀ 0.0001 in LB medium supplemented with 100 mg/L Tet, 50 mg/L Gm, and 200 µM cumate, followed by incubation at 37°C, 170 rpm for 12 hours. After 4 hours of incubation, 200 µM additional cumate is added to the cultures. After incubation, 3 µL of culture was placed on a 1% agarose pad with or without 100 mg/L Tet, 50 mg/L Gm, and 200 µM cumate, then imaged under the same conditions. For fluorescent imaging of WT and *mreB* strain (PAMNT054 with sfGFP expression based on plasmid). Lattice SIM imaging was performed using a Zeiss Elyra 7 AxioObserver inverted microscope equipped with 488 nm (100 mW), 561 nm (100 mW), and 642 nm (150 mW) laser lines, operated at 20% of their maximum output. A Plan-Apochromat 63×/1.4 NA Oil DIC M27 objective (Zeiss) and a 1.6× magnification lens in the detection path were used, resulting in a raw image pixel size of 64.5 nm. Fluorescence emission was separated from excitation using a dichroic beamsplitter (405/488/561/642) and a dual-band emission filter (495–550 nm and 570–620 nm). Images were captured using a PCO Edge 4.2 sCMOS camera, and acquisition was controlled via Zen Black software (Zeiss). The Lattice SIM technique used a patterned point illumination and laterally shifted phase images, with 13 phase images captured per plane per channel. Long-time-lapse acquisitions were performed using the epifluorescence technique. Exposure settings per phase were 100 ms at 1% laser power for the 488 nm line and 50 ms at 1% for the 642 nm line. The sample temperature was maintained at 37 °C during imaging. Time-lapse acquisitions consisted of either 41 cycles with 5-minute intervals between frames. Illumination grids of 27.5 µm (488 nm) and 32 µm (642 nm) were selected to optimize modulation contrast. Image reconstruction was performed using Zen Black’s nonlinear iterative SIM algorithm with 2D+ processing enabled. General settings were “Live” and “Weak,” and advanced filters were set to “Best fit” and “Median,” with a sectioning value of 100 and baseline subtraction enabled. Final reconstructed images had a lateral pixel size of 16.1 nm. Cell segmentation was performed either manually using the software Napari^89^ or semi-automatically. The semi-automatic method consisted of first performing deep-learning-based segmentation with Omnipose, either with a pre-trained model for WT-looking cells or with a custom-trained model for spherical cells. Cell morphology (width, length) was extracted by measuring the bounding rectangle of the corresponding cell mask. Growth curves were obtained as the number of cells detected per frame, normalised to the number in the first frame. Doubling times were extracted by comparing the number of cells in the first and last frame of the experiment, following the equation:

τ = Δt.*ln*(2)/*ln(N_end_/N_beginning_)*, where Δt is the time between the beginning and the end of the experiment, N_beginning_ is the number of cells in the first frame, and N_end_ is the number of cells in the last frame.

The cell circularity level and cell volume of both WT and Δ*mreB* were calculated according to these equations:

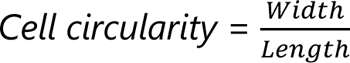

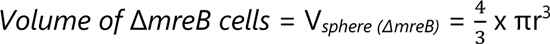, where r is the average width of the cell.

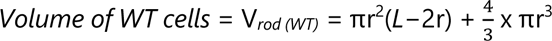, where L is the length of the cells and r is the width.

### Swarming motility assay

A swarming motility assay was performed using a nutrient-rich, low-agar medium to facilitate bacterial movement. The medium consisted of 0.5% (w/v) agar, 8 g/L nutrient broth, and distilled water, which was autoclaved to ensure sterility. After autoclaving, glucose was added to a final concentration of 0.5% (w/v) and mixed thoroughly. The medium was allowed to cool and dry for at least 30 minutes at RT without stacking to ensure a uniform surface. Overnight bacterial cultures (ON cultures) were grown in LB at 37°C, 170 rpm. To complete swarming assays, 2 µL of ON culture was carefully spotted onto the centre of each plate. The inoculated plates were incubated at 30°C for 16 hours to allow swarming motility.

### Twitching motility assay

The macroscopic twitching assay protocol of Turnbull et al., 2014 is used^91^. 1% LB Agar solid plates were prepared. The solid cultures were streaked on 1.5% LB agar plates at 37°C for ON. Using a plastic inoculation loop, a small portion of the outer edge of the bacterial streak was taken and mixed into a sterile area of agar until the culture was smooth. Then a match-head-sized inoculum from the mixed culture was placed in the middle of a 1% LB agar plate using a pipette tip, ensuring the tip touched the edge of the plate. Plates were incubated at 37°C for 48 hours and were imaged under white light to observe the halo structure of bacteria that have twitched across the plate between the bottom of the agar and the plastic petri plate (interstitial colony). Growth of colonies was observed and analysed using a fully automated, AI-powered petri and microtiter plate analysis machine from Reshape Biotech (Denmark).

### Swimming mobility assay

The swimming assay protocol of Ha et al., 2022, is optimised for our conditions^92^. 0.25% LB Agar solid plates were prepared. The medium was allowed to cool and dry for at least 30 minutes at RT without stacking to ensure a uniform surface. Overnight bacterial cultures (ON cultures) were grown in LB at 37°C, 170 rpm. To complete swimming assays, 5 µL of ON culture (OD_600_ nm was adjusted to 1.0 for all strains) was carefully spotted onto the centre of each plate. The inoculated plates were incubated at 37°C for 24 hours to allow swimming motility. Images of swimming were taken under white light, and growth was observed and analysed by a fully automated AI-powered petri and microtiter plate analysis machine, Reshape Biotech (Denmark).

### Scanning electron microscopy

*P. aeruginosa* strains were grown in LB medium to the exponential phase (OD_600_nm 0.15–0.20), then 50 µL of culture of each strain was fixed separately in 2 mL of 2% glutaraldehyde buffered with 0.2 M sodium cacodylate (EMS 12300, LFG France) for 2 hours at RT, followed by overnight incubation at 4°C. Fixation was performed in a 24-well plate (TPP 92024, Switzerland) containing 1 cm × 1 cm glass slides, which were pre-cleaned with ethanol, plasma-treated, sonicated, and coated with 0.1 M Poly-L-Lysine (Sigma Merck P8920). The attached cells were rinsed twice in 0.2 M sodium cacodylate buffer, dehydrated through ethanol baths (50%, 70%, 90%, 100%, and anhydrous 100%), and dried using a Leica EM300 critical point apparatus with 20 slow exchange cycles. Samples were mounted on aluminium stubs with adhesive carbon (EMS, LFG France) and coated with a 6 nm Au/Pd layer using a Quorum SC7620 sputter coater with 50 Pa of Ar, 180 s of sputtering at 3.5 mA. Imaging was conducted with a SE detector of a Hitachi SU5000 FEG-SEM at 2 keV, 30 spot size, and 5 mm working distance at the MIMA2 core facility, INRAE, Jouy-en-Josas, France; https://doi.org/10.15454/1.5572348210007727E12.

### Transmission electron microscopy

The same exponential-phase cultures used for SEM imaging were fixed in 2% glutaraldehyde prepared in 0.1 M sodium cacodylate buffer (pH 7.2) for 4 hours at room temperature to preserve cellular structures. Following fixation, samples were contrasted with 0.2% Oolong Tea Extract (OTE) in cacodylate buffer and post-fixed with 1% osmium tetroxide containing 1.5% potassium ferrocyanide to enhance membrane contrast. Dehydration was performed through a graded ethanol series (30% to 100%), followed by gradual substitution with an ethanol-Epon mixture before embedding in Epon resin (Delta Microscopie, France). Thin sections (70 nm) were collected on 200-mesh copper grids and counterstained with lead citrate to improve contrast. Imaging was conducted using a Hitachi HT7700 transmission electron microscope (Milexia, France) operated at 80 kV, and images were captured with a charge-coupled device (CCD) camera (AMT). This study utilized the facilities and expertise of the MIMA2 Microscopy and Imaging Facility for Microbes, Animals and Foods https://doi.org/10.15454/1.5572348210007727E12.

### MIC determination

Minimum Inhibitory Concentrations of WT and *mreB* mutant (PAMNT054) strains were quantified using the EUCAST broth microdilution reference method, as recommended for MIC determination. Pre-cultures of strains were prepared in Mueller-Hinton Broth (MH) medium with 0.5 McFarland (≈10^8^ CFU/mL). These cultures were inoculated into ready-to-use antibiotic plates (Sensititre™ EU Surveillance EUVSEC3 AST, Thermo Fisher Scientific, East Grinstead, UK) to determine MIC values. They were incubated at 37°C for 24 hours with agitation, and MIC values were determined at the end of 24 hours.

### Biofilm formation and analysis

Fluorescently labelled strains were grown overnight in LB medium supplemented with 50 mg/L Gm to maintain plasmids. However, no antibiotics were added during the biofilm formation step to avoid interfering with the biofilm structure. To confirm the stability of the plasmids in biofilms without antibiotic exposure, separate stability assays were performed, and no significant differences in plasmid retention were observed (Fig. S16). 200 µL monoculture (OD_600_nm 0.05), 3:1, 3:2, 1:1, 2:3, and 1:3 (WT:Δ*mreB*) ratio co-cultures were prepared in polystyrene 96-well microtiter plates with a µclear® base (Greiner Bio-One, France), adapted for high-resolution fluorescence imaging. After 2 hours at 37°C, the media (LB) of the biofilm was refreshed to discard non-adherent bacteria. Then, the microtiter plate was incubated under a microscope for 16 h at 37°C; images were taken every 1.5 h to monitor biofilm development. 3D images were acquired by SP8 Leica Laser Scanning Confocal Microscopy by HCS method with 1 µm z-step, 2 lines average, 1 line accumulation, 512×512 area, 120 frames, with a water 63 × immersion lens (NA = 1.2), excitation wavelength of 488 nm 5% and 561 nm 5% with emission wavelengths collected from 500 to 550 nm and from 590 to 700 nm respectively for sfGFP and mScarletI fluorescence. Three-dimensional projections of biofilm structures were reconstructed using the Easy 3D function in IMARIS 10.1 (Bitplane, Switzerland). Biofilm biovolumes (µm^3^) and rugosity extracted from CLSM images were analysed with BiofilmQ software^93^.

### Flow cytometry

Overnight cultures of WT, Δ*mreB* and *mreBCD, mreCD* silenced strains were diluted to OD_600_nm 0.01 in 100 mL LB medium supplemented with 100 mg/L Tet and 50 mg/L Gm, and when OD_600_nm reached 0.15-0.20, the culture was split into 2 subcultures and 200 µM cumate was added to only one of them. After culturing, 300 µL of culture was collected from each condition and fixed with 0.1 M KPO4, 16% PFA, and 25% Glutaraldehyde at 3-, 5-, and 7-hour time points. Fixed samples were observed by Flowcytometry (Beckman Coulter’s benchtop CytoFLEX flow cytometer). Acquisition settings are listed in (Fig. S8a), and instead of all events, only the *P. aeruginosa* gate is created by fluorescent labelling, and this gate is used for event counting in WT and shifting from this area in mutant cells added in each condition separately. To observe event count differences in culture over time, 5 µL of a 10x-diluted culture in 0.85% NaCl dH_2_O was spread.

## Data Availability

Relevant research data will be deposited on Zenodo after publication and are available from the corresponding authors upon request. Constructs can be available from the corresponding authors under a material transfer agreement. The Whole Genome Shotgun project has been deposited at DDBJ/ENA/GenBank under the accession numbers <u>JBTKTW000000000</u>, <u>JBTKTV000000000</u>, <u>JBTKTU000000000</u>, <u>JBTKTT000000000</u>, <u>JBTKTS000000000</u>, <u>JBTKTR000000000</u> for strains PAMNT043, PAMNT044, PAMNT048, PAMNT049, PAMNT051 and PAMNT054 respectively and the draft genome assembly and annotation can be found in NCBI under BioProject number <u>PRJNA1265664</u> and SRA numbers, <u>SRR36779680</u>, <u>SRR36779679</u>, <u>SRR36779678</u>, <u>SRR36779677</u>, <u>SRR36779676</u>, <u>SRR36779675</u> for PAMNT043, PAMNT044, PAMNT048, PAMNT049, PAMNT051 and PAMNT054 respectively.

## Acknowledgements

We thank Craig MacLean for plasmids pSEVA627-empty_sfgfp and pSEVA627M-empty_mScarletI, and Ina Attree for pJN105 plasmid with *mreC, mreD* and *mreCD* extra copies, the anti-MreC antibodies and western blot protocol for FtsZ. We thank Pierre Renaud for his sequencing and bioinformatics expertise, and the members of the ProCeD and B3D teams for helpful discussions. This work has benefited from the equipment and expertise of MIMA2 Imaging Core Facility (Université Paris-Saclay, INRAE, AgroParisTech, 78350, Jouy-en-Josas, France, DOI: https://doi.org/10.15454/1.5572348210007727E12) for TEM and SEM acquisitions; we thank Christine Péchoux for the TEM acquisitions and Vlad Costache for the SEM acquisitions. This work was funded by a grant from the European Research Council (ERC) under the Horizon 2020 research and innovation program (ERC CoG No 772178 to R.C.-L.). M.-N.T.’s PhD was funded by a PhD fellowship from the Region Ile-de-France (collaborative project DIM1HEALTH, nr R21157DP to R.B. and R.C.-L.) and supported by a travel grant Erasmus+.

## References

1. Hugonneau-Beaufet, I. et al. Characterization of Pseudomonas aeruginosa L,D - Transpeptidases and Evaluation of Their Role in Peptidoglycan Adaptation to Biofilm Growth. Microbiol Spectr 11, e05217–22 (2023).

2. Thi, M. T. T., Wibowo, D. & Rehm, B. H. A. Pseudomonas aeruginosa Biofilms. IJMS 21, 8671 (2020).

3. Martinez, J. L. et al. A global view of antibiotic resistance. FEMS Microbiol Rev 33, 44–65 (2009).

4. Vollmer, W., Blanot, D. & De Pedro, M. A. Peptidoglycan structure and architecture. FEMS Microbiol Rev 32, 149–167 (2008).

5. Typas, A., Banzhaf, M., Gross, C. A. & Vollmer, W. From the regulation of peptidoglycan synthesis to bacterial growth and morphology. Nat Rev Microbiol 10, 123–136 (2012).

6. Den Blaauwen, T., De Pedro, M. A., Nguyen-Distèche, M. & Ayala, J. A. Morphogenesis of rod-shaped sacculi. FEMS Microbiol Rev 32, 321–344 (2008).

7. Egan, A. J. F., Errington, J. & Vollmer, W. Regulation of peptidoglycan synthesis and remodelling. Nat Rev Microbiol 18, 446–460 (2020).

8. Zapun, A., Contreras-Martel, C. & Vernet, T. Penicillin-binding proteins and β-lactam resistance. FEMS Microbiol Rev 32, 361–385 (2008).

9. Caveney, N. A. et al. Structure of the Peptidoglycan Synthase Activator LpoP in Pseudomonas aeruginosa. Structure 28, 643–650.e5 (2020).

10. Cho, H. et al. Bacterial cell wall biogenesis is mediated by SEDS and PBP polymerase families functioning semi-autonomously. Nat Microbiol 1, 16172 (2016).

11. Chen, W., Zhang, Y.-M. & Davies, C. Penicillin-Binding Protein 3 Is Essential for Growth of Pseudomonas aeruginosa. Antimicrob Agents Chemother 61, e01651–16 (2017).

12. Vigouroux, A. et al. Class-A penicillin binding proteins do not contribute to cell shape but repair cell-wall defects. eLife 9, e51998 (2020).

13. Shi, H., Bratton, B. P., Gitai, Z. & Huang, K. C. How to Build a Bacterial Cell: MreB as the Foreman of E. coli Construction. Cell 172, 1294–1305 (2018).

14. Domínguez-Escobar, J. et al. Processive Movement of MreB-Associated Cell Wall Biosynthetic Complexes in Bacteria. Science 333, 225–228 (2011).

15. Garner, E. C. et al. Coupled, Circumferential Motions of the Cell Wall Synthesis Machinery and MreB Filaments in B. subtilis. Science 333, 222–225 (2011).

16. Carballido-Lopez, R. The actin-like MreB proteins in Bacillus subtilis a new turn. Front Biosci **S**4, 1582–1606 (2012).

17. Jones, L. J. F., Carballido-López, R. & Errington, J. Control of Cell Shape in Bacteria. Cell 104, 913–922 (2001).

18. Bendezú, F. O., Hale, C. A., Bernhardt, T. G. & De Boer, P. A. J. RodZ (YfgA) is required for proper assembly of the MreB actin cytoskeleton and cell shape in E. coli. EMBO J 28, 193–204 (2009).

19. Robertson, G. T. et al. A Novel Indole Compound That Inhibits *Pseudomonas aeruginosa* Growth by Targeting MreB Is a Substrate for MexAB-OprM. J Bacteriol 189, 6870–6881 (2007).

20. Spiers, A. J., Kahn, S. G., Bohannon, J., Travisano, M. & Rainey, P. B. Adaptive Divergence in Experimental Populations of *Pseudomonas fluorescens*. I. Genetic and Phenotypic Bases of Wrinkly Spreader Fitness. Genetics 161, 33–46 (2002).

21. Noirot-Gros, M.-F., Forrester, S., Malato, G., Larsen, P. E. & Noirot, P. CRISPR interference to interrogate genes that control biofilm formation in Pseudomonas fluorescens. Sci Rep 9, 15954 (2019).

22. Bean, G. J. et al. A22 Disrupts the Bacterial Actin Cytoskeleton by Directly Binding and Inducing a Low-Affinity State in MreB. Biochemistry 48, 4852–4857 (2009).

23. Yamachika, S. et al. Anti-Pseudomonas aeruginosa Compound, 1,2,3,4-Tetrahydro-1,3,5-triazine Derivative, Exerts Its Action by Primarily Targeting MreB. Biological & Pharmaceutical Bulletin 35, 1740–1744 (2012).

24. Cowles, K. N. & Gitai, Z. Surface association and the MreB cytoskeleton regulate pilus production, localization and function in Pseudomonas aeruginosa: Pseudomonas pilus regulation and MreB. Molecular Microbiology 76, 1411–1426 (2010).

25. Bonez, P. C. et al. Anti-biofilm activity of A22 ((S-3,4-dichlorobenzyl) isothiourea hydrochloride) against Pseudomonas aeruginosa: Influence on biofilm formation, motility and bioadhesion. Microbial Pathogenesis 111, 6–13 (2017).

26. Smith, W. P. J. et al. Cell morphology drives spatial patterning in microbial communities. Proc. Natl. Acad. Sci. U.S.A. 114, (2017).

27. Pazos, M. & Peters, K. Peptidoglycan. in Bacterial Cell Walls and Membranes (ed. Kuhn, A.) vol. 92 127–168 (Springer International Publishing, Cham, 2019).

28. Martins, A. et al. Self-association of MreC as a regulatory signal in bacterial cell wall elongation. Nat Commun 12, 2987 (2021).

29. Rohs, P. D. A. et al. A central role for PBP2 in the activation of peptidoglycan polymerization by the bacterial cell elongation machinery. PLoS Genet 14, e1007726 (2018).

30. Contreras-Martel, C. et al. Molecular architecture of the PBP2–MreC core bacterial cell wall synthesis complex. Nat Commun 8, 776 (2017).

31. Valentin, J. D. P. et al. Identification of Potential Antimicrobial Targets of Pseudomonas aeruginosa Biofilms through a Novel Screening Approach. Microbiol Spectr 11, e03099–22 (2023).

32. Wang, S., Furchtgott, L., Huang, K. C. & Shaevitz, J. W. Helical insertion of peptidoglycan produces chiral ordering of the bacterial cell wall. Proc. Natl. Acad. Sci. U.S.A. 109, (2012).

33. White, C. L., Kitich, A. & Gober, J. W. Positioning cell wall synthetic complexes by the bacterial morphogenetic proteins MreB and MreD: Control of cell shape in bacteria. Molecular Microbiology 76, 616–633 (2010).

34. Kruse, T., Bork-Jensen, J. & Gerdes, K. The morphogenetic MreBCD proteins of *Escherichia coli* form an essential membrane-bound complex. Molecular Microbiology 55, 78–89 (2005).

35. Bendezú, F. O. & De Boer, P. A. J. Conditional Lethality, Division Defects, Membrane Involution, and Endocytosis in mre and mrd Shape Mutants of Escherichia coli. J Bacteriol 190, 1792–1811 (2008).

36. Divakaruni, A. V., Loo, R. R. O., Xie, Y., Loo, J. A. & Gober, J. W. The cell-shape protein MreC interacts with extracytoplasmic proteins including cell wall assembly complexes in *Caulobacter crescentus*. Proc. Natl. Acad. Sci. U.S.A. 102, 18602–18607 (2005).

37. Tavares, A. C., Fernandes, P. B., Carballido-López, R. & Pinho, M. G. MreC and MreD Proteins Are Not Required for Growth of Staphylococcus aureus. PLoS ONE 10, e0140523 (2015).

38. Land, A. D. & Winkler, M. E. The Requirement for Pneumococcal MreC and MreD Is Relieved by Inactivation of the Gene Encoding PBP1a. J Bacteriol 193, 4166–4179 (2011).

39. Yulo, P. R. J. et al. Evolutionary rescue of spherical mreB deletion mutants of the rod-shape bacterium Pseudomonas fluorescens SBW25. eLife 13, RP98218 (2025).

40. Kruse, T. Dysfunctional MreB inhibits chromosome segregation in Escherichia coli. The EMBO Journal 22, 5283–5292 (2003).

41. Nilsen, T., Yan, A. W., Gale, G. & Goldberg, M. B. Presence of Multiple Sites Containing Polar Material in Spherical*Escherichia coli*Cells That Lack MreB. J Bacteriol 187, 6187–6196 (2005).

42. Wachi, M. et al. Mutant isolation and molecular cloning of mre genes, which determine cell shape, sensitivity to mecillinam, and amount of penicillin-binding proteins in Escherichia coli. J Bacteriol 169, 4935–4940 (1987).

43. Tadić, V., Josipović, G., Zoldoš, V. & Vojta, A. CRISPR/Cas9-based epigenome editing: An overview of dCas9-based tools with special emphasis on off-target activity. Methods 164–165, 109–119 (2019).

44. Qi, L. S. et al. Repurposing CRISPR as an RNA-Guided Platform for Sequence-Specific Control of Gene Expression. Cell 152, 1173–1183 (2013).

45. Gray, D. A. et al. Extreme slow growth as alternative strategy to survive deep starvation in bacteria. Nat Commun 10, 890 (2019).

46. Shimaya, T., Okura, R., Wakamoto, Y. & Takeuchi, K. A. Scale invariance of cell size fluctuations in starving bacteria. Commun Phys 4, 238 (2021).

47. Liu, Z., Zhao, Q., Xu, C. & Song, H. Compensatory evolution of chromosomes and plasmids counteracts the plasmid fitness cost. Ecology and Evolution 14, e70121 (2024).

48. Pinho, M. G., Kjos, M. & Veening, J.-W. How to get (a)round: mechanisms controlling growth and division of coccoid bacteria. Nat Rev Microbiol 11, 601–614 (2013).

49. Köhler, T., Curty, L. K., Barja, F., Van Delden, C. & Pechère, J.-C. Swarming of *Pseudomonas aeruginosa* Is Dependent on Cell-to-Cell Signaling and Requires Flagella and Pili. J Bacteriol 182, 5990–5996 (2000).

50. Zegadło, K. et al. Bacterial Motility and Its Role in Skin and Wound Infections. IJMS 24, 1707 (2023).

51. Cowles, K. N. & Gitai, Z. Surface association and the MreB cytoskeleton regulate pilus production, localization and function in Pseudomonas aeruginosa: Pseudomonas pilus regulation and MreB. Molecular Microbiology 76, 1411–1426 (2010).

52. Caiazza, N. C., Shanks, R. M. Q. & O’Toole, G. A. Rhamnolipids Modulate Swarming Motility Patterns of *Pseudomonas aeruginosa*. J Bacteriol 187, 7351–7361 (2005).

53. Yeung, A. T. Y. et al. Swarming of *Pseudomonas aeruginosa* Is Controlled by a Broad Spectrum of Transcriptional Regulators, Including MetR. J Bacteriol 191, 5592–5602 (2009).

54. Mauriello, E. M. F. et al. Bacterial motility complexes require the actin-like protein, MreB and the Ras homologue, MglA. EMBO J 29, 315–326 (2010).

55. Francis, V. I. et al. Multiple communication mechanisms between sensor kinases are crucial for virulence in Pseudomonas aeruginosa. Nat Commun 9, (2018).

56. Deforet, M., Van Ditmarsch, D., Carmona-Fontaine, C. & Xavier, J. B. Hyperswarming adaptations in a bacterium improve collective motility without enhancing single cell motility. Soft Matter 10, 2405–2413 (2014).

57. Morita, Y., Tomida, J. & Kawamura, Y. Resistance and Response to Anti-Pseudomonas Agents and Biocides. 173–187 (2015) doi:10.1007/978-94-017-9555-5_7.

58. Fàbrega, A., Madurga, S., Giralt, E. & Vila, J. Mechanism of action of and resistance to quinolones. Microbial Biotechnology 2, 40–61 (2009).

59. Shariati, A. et al. The resistance mechanisms of bacteria against ciprofloxacin and new approaches for enhancing the efficacy of this antibiotic. Front. Public Health 10, 1025633 (2022).

60. Ranjitkar, S. et al. Target (MexB)- and Efflux-Based Mechanisms Decreasing the Effectiveness of the Efflux Pump Inhibitor D13-9001 in *Pseudomonas aeruginosa* PAO1: Uncovering a New Role for MexMN-OprM in Efflux of β-Lactams and a Novel Regulatory Circuit (MmnRS) Controlling MexMN Expression. Antimicrob Agents Chemother 63, e01718–18 (2019).

61. Piddock, L. J. V., Jin, Y.-F., Ricci, V. & Asuquo, A. E. Quinolone accumulation by Pseudomonas aeruginosa, Staphylococcus aureus and Escherichia coli. Journal of Antimicrobial Chemotherapy 43, 61–70 (1999).

62. Strahl, H., Bürmann, F. & Hamoen, L. W. The actin homologue MreB organizes the bacterial cell membrane. Nat Commun 5, 3442 (2014).

63. Legaree, B. A., Daniels, K., Weadge, J. T., Cockburn, D. & Clarke, A. J. Function of penicillin-binding protein 2 in viability and morphology of Pseudomonas aeruginosa. Journal of Antimicrobial Chemotherapy 59, 411–424 (2007).

64. Montaner, M., Lopez-Argüello, S., Oliver, A. & Moya, B. PBP Target Profiling by β-Lactam and β-Lactamase Inhibitors in Intact Pseudomonas aeruginosa: Effects of the Intrinsic and Acquired Resistance Determinants on the Periplasmic Drug Availability. Microbiol Spectr 11, e03038–22 (2023).

65. Kocaoglu, O. & Carlson, E. E. Profiling of β-Lactam Selectivity for Penicillin-Binding Proteins in Escherichia coli Strain DC2. Antimicrob Agents Chemother 59, 2785–2790 (2015).

66. Georgopapadakou, N. H., Dix, B. A. & Mauriz, Y. R. Possible physiological functions of penicillin-binding proteins in Staphylococcus aureus. Antimicrob Agents Chemother 29, 333–336 (1986).

67. Glen, K. A. & Lamont, I. L. Penicillin-binding protein 3 sequence variations reduce susceptibility of *Pseudomonas aeruginosa* to β-lactams but inhibit cell division. Journal of Antimicrobial Chemotherapy 79, 2170–2178 (2024).

68. Dion, M. F. et al. Bacillus subtilis cell diameter is determined by the opposing actions of two distinct cell wall synthetic systems. Nat Microbiol 4, 1294–1305 (2019).

69. Veyrier, F. J. et al. Common Cell Shape Evolution of Two Nasopharyngeal Pathogens. PLoS Genet 11, e1005338 (2015).

70. Vollmer, W. The prokaryotic cytoskeleton: a putative target for inhibitors and antibiotics? Appl Microbiol Biotechnol 73, 37–47 (2006).

71. Harshey, R. M. Bacterial Motility on a Surface: Many Ways to a Common Goal. Annu. Rev. Microbiol. 57, 249–273 (2003).

72. Murga, R., Stewart, P. S. & Daly, D. Quantitative analysis of biofilm thickness variability. Biotech & Bioengineering 45, 503–510 (1995).

73. Oliveira, M., Cunha, E., Tavares, L. & Serrano, I. *P. aeruginosa* interactions with other microbes in biofilms during co-infection. AIMSMICRO 9, 612–646 (2023).

74. Monmeyran, A. et al. The inducible chemical-genetic fluorescent marker FAST outperforms classical fluorescent proteins in the quantitative reporting of bacterial biofilm dynamics. Sci Rep 8, 10336 (2018).

75. Stackebrandt, E. & Woese, C. R. A phylogenetic dissection of the family micrococcaceae. Current Microbiology 2, 317–322 (1979).

76. Siefert’t, J. L. & Fox, G. E. Phylogenetic mapping of bacterial morphology. Microbiology 144, 2803–2808 (1998).

77. Diaz Caballero, J., et al. Selective Sweeps and Parallel Pathoadaptation Drive Pseudomonas aeruginosa Evolution in the Cystic Fibrosis Lung. mBio 6, e00981–15 (2015).

78. Pas, E. Perpendicular planes of FtsZ arcs in spheroidal Escherichia coli cells. Biochimie 83, 121–124 (2001).

79. Erickson, H. P., Anderson, D. E. & Osawa, M. FtsZ in Bacterial Cytokinesis: Cytoskeleton and Force Generator All in One. Microbiol Mol Biol Rev 74, 504–528 (2010).

80. Valentin, J. D. P. et al. Role of the flagellar hook in the structural development and antibiotic tolerance of Pseudomonas aeruginosa biofilms. ISME J 16, 1176–1186 (2022).

81. Robertson, G. T. et al. A Novel Indole Compound That Inhibits *Pseudomonas aeruginosa* Growth by Targeting MreB Is a Substrate for MexAB-OprM. J Bacteriol 189, 6870–6881 (2007).

82. Bryan, E. J. et al. TXH11106: A Third-Generation MreB Inhibitor with Enhanced Activity against a Broad Range of Gram-Negative Bacterial Pathogens. Antibiotics 11, 693 (2022).

83. Hmelo, L. R. et al. Precision-engineering the Pseudomonas aeruginosa genome with two-step allelic exchange. Nat Protoc 10, 1820–1841 (2015).

84. Topp, S. et al. Synthetic Riboswitches That Induce Gene Expression in Diverse Bacterial Species. Appl Environ Microbiol 76, 7881–7884 (2010).

85. Cui, W. et al. Engineering an inducible gene expression system for Bacillus subtilis from a strong constitutive promoter and a theophylline-activated synthetic riboswitch. Microb Cell Fact 15, 199 (2016).

86. Seo, S.-O. & Schmidt-Dannert, C. Development of a synthetic cumate-inducible gene expression system for Bacillus. Appl Microbiol Biotechnol 103, 303–313 (2019).

87. Tunç, M. N. et al. Genome sequences of four colistin-resistant ESKAPE bacterial strains isolated from patients within the same hospital. Microbiol Resour Announc 13, e00874–23 (2024).

88. Clausen, P. T. L. C., Aarestrup, F. M. & Lund, O. Rapid and precise alignment of raw reads against redundant databases with KMA. BMC Bioinformatics 19, 307 (2018).

89. Chiu, C.-L., Clack, N., & the napari community. napari: a Python Multi-Dimensional Image Viewer Platform for the Research Community. Microscopy and Microanalysis 28, 1576–1577 (2022).

90. Cutler, K. J. et al. Omnipose: a high-precision morphology-independent solution for bacterial cell segmentation. Nat Methods 19, 1438–1448 (2022).

91. Turnbull, L. & Whitchurch, C. B. Motility Assay: Twitching Motility. in Pseudomonas Methods and Protocols (eds Filloux, A. & Ramos, J.-L.) vol. 1149 73–86 (Springer New York, New York, NY, 2014).

92. Ha, D.-G., Kuchma, S. L. & O’Toole, G. A. Plate-Based Assay for Swimming Motility in Pseudomonas aeruginosa. in Pseudomonas Methods and Protocols (eds Filloux, A. & Ramos, J.-L.) vol. 1149 59–65 (Springer New York, New York, NY, 2014).

93. Hartmann, R. et al. Quantitative image analysis of microbial communities with BiofilmQ. Nat Microbiol 6, 151–156 (2021).

