## Supplementary Figures for "MreB is dispensable for viability but critical for rod shape, motility and biofilm fitness in *Pseudomonas aeruginosa*"

### Fig. S1

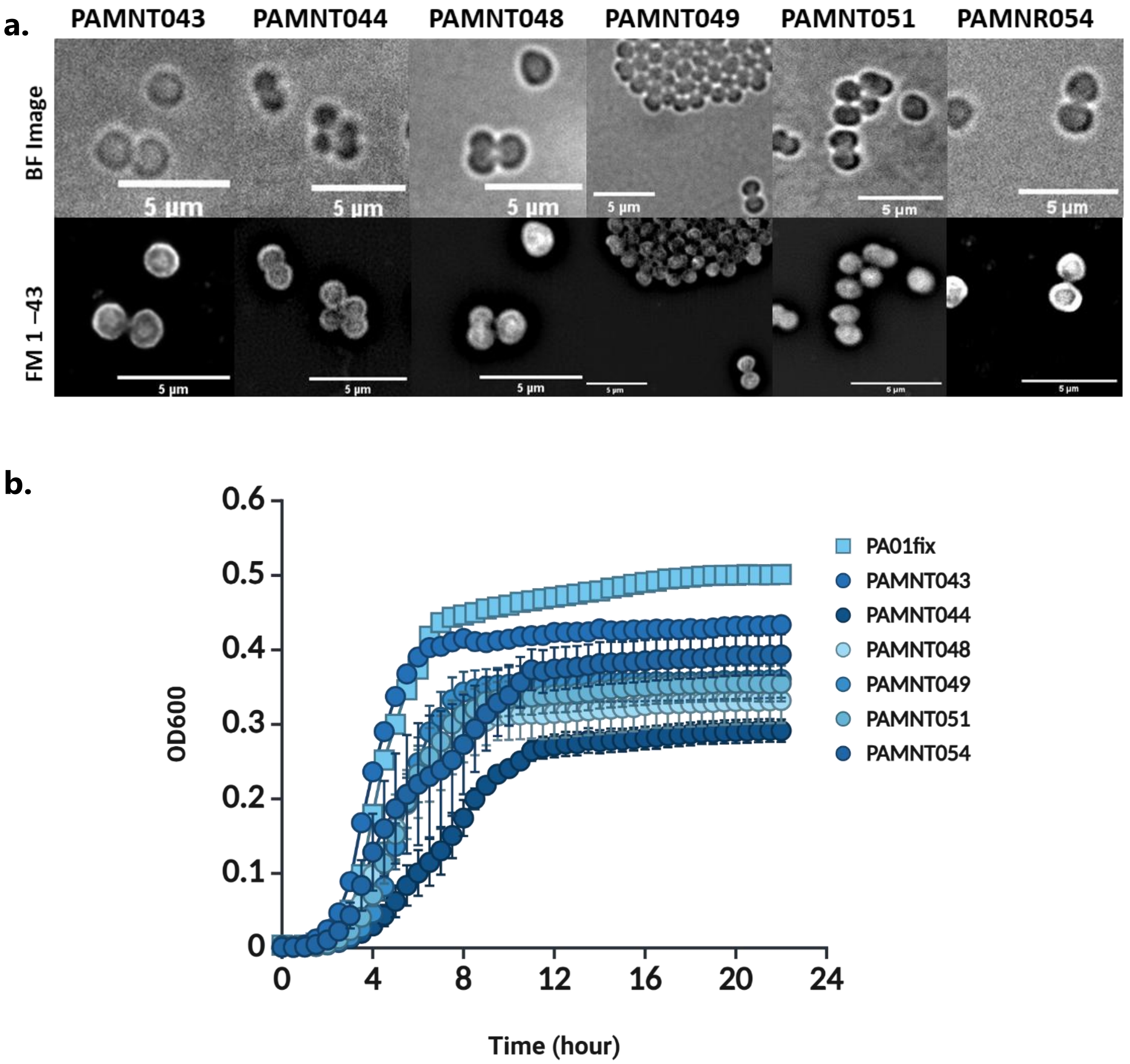


**Fig. S1: (a)** BF and FM1-43 membrane labelled images of fixed *mreB* deleted strains. All 6 strains showed spherical morphology. Scale bar is 5 µm. **(b)** Growth curves of *mreB* deleted strains, parental strain (PA01-fix). There are no significant changes in duplication time within mreB-deleted strains.

**Fig. S2**

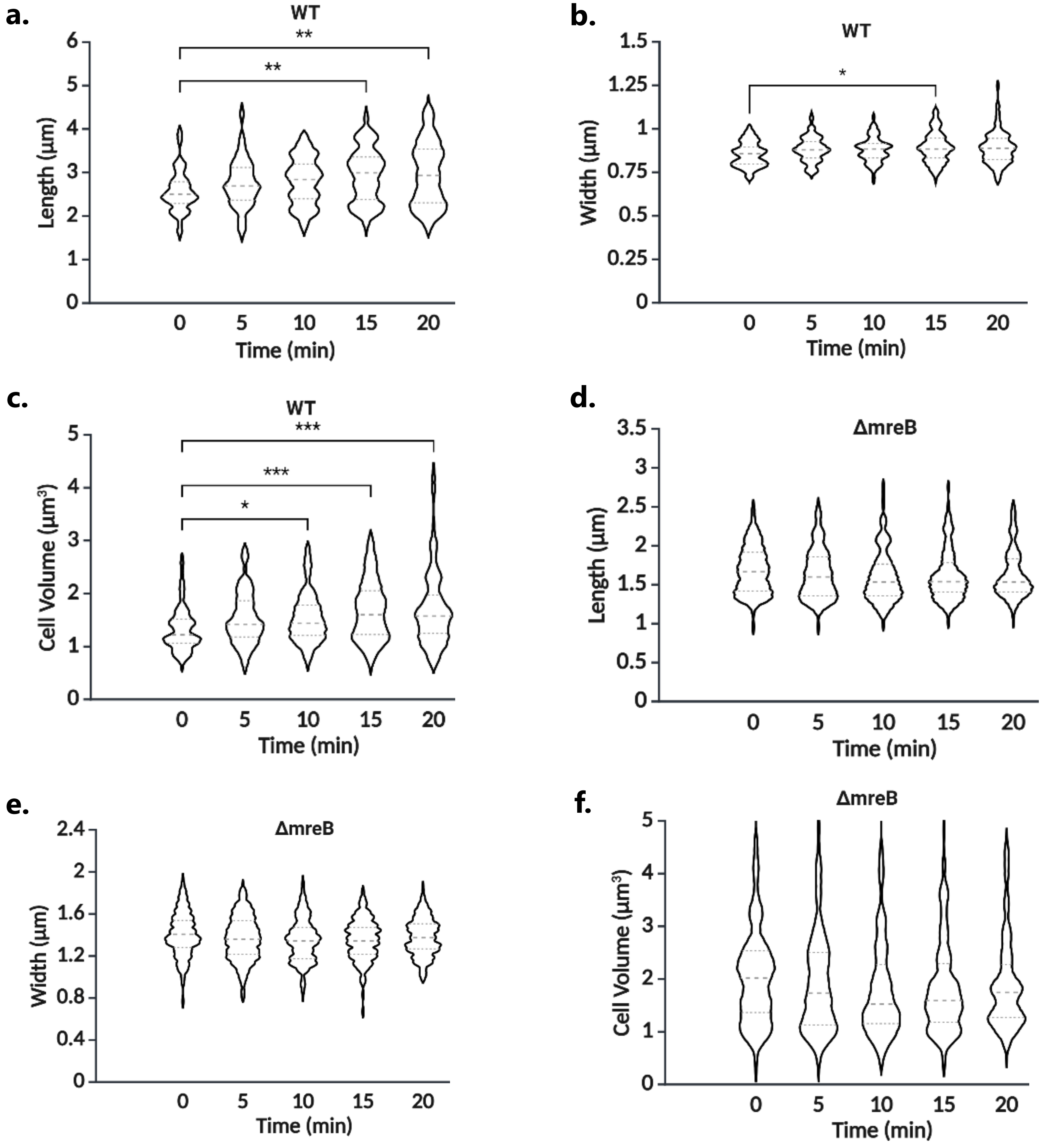


**Fig. S2a, b, c, d, e, f:** Length **(a)**, width **(b)** and cell volume **(c)** measurement of WT cells over 20 minutes with 5 min intervals, starting from the first frame after the cells are put on an LB 1% agarose pad. Kruskal-Wallis test with Dunn's multiple comparisons test was applied separately. The initial time point (0) was found to be significantly lower than several later time points: for length, $p = 0.0019$ (0 vs. 15 min) and $p = 0.0011$ (0 vs. 20 min); for width, $p = 0.0236$ (0 vs. 15 min); for cell volume, $p = 0.0356$ (0 vs. 10 min), $p = 0.0002$ (0 vs. 15 min) and $p = 0.0004$ (0 vs. 20 min). Length **(d)**, width **(e)** and cell volume **(f)** measurement of *ΔmreB* cells over 20 minutes with 5 min intervals. 0 is the first frame after the cells are put on an LB 1% agarose pad. Kruskal-Wallis test with Dunn's multiple comparisons test was applied separately, and no differences were found.

#### Fig. S3

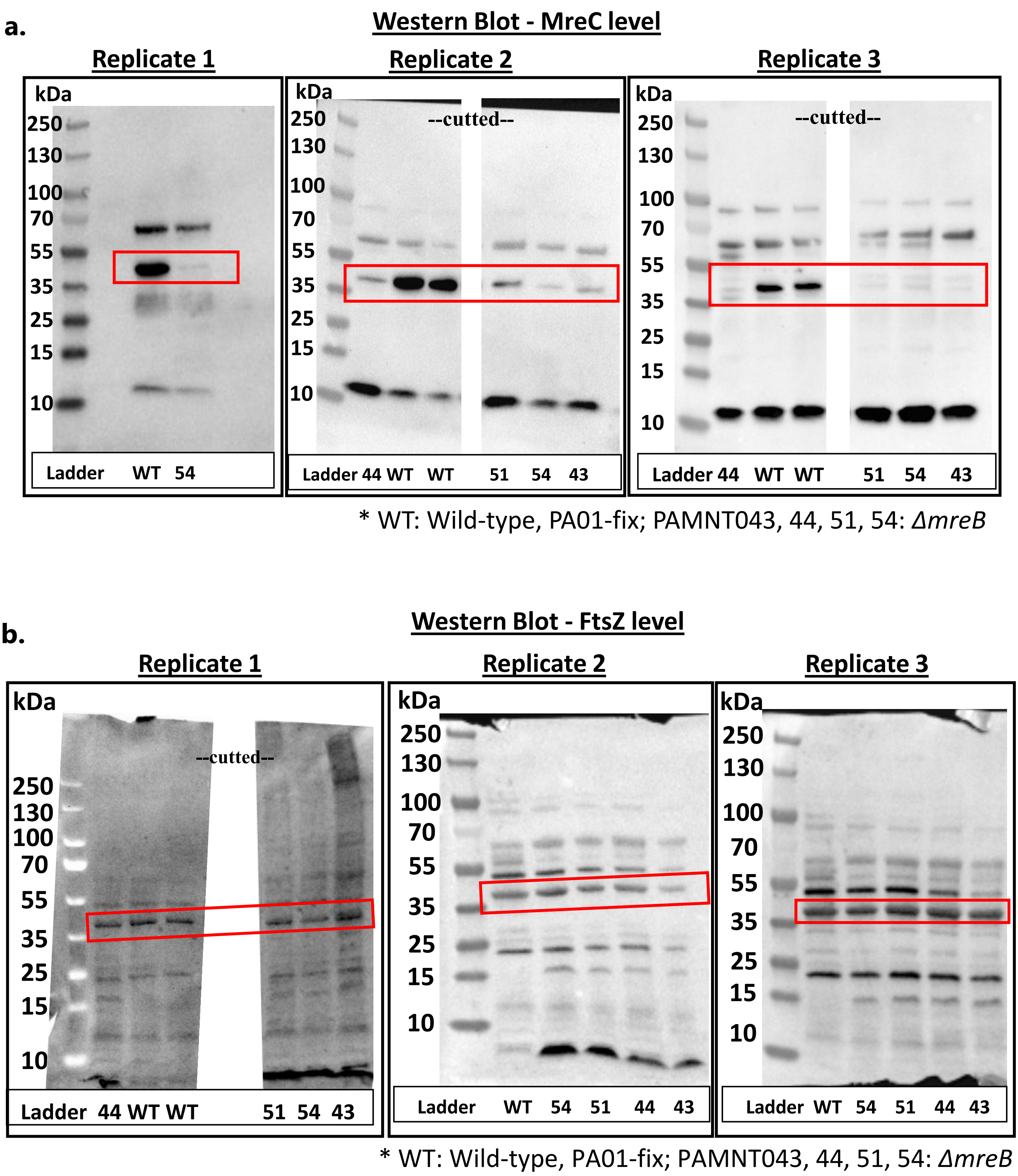


**Fig. S3: (a)** Western blot analysis of MreC levels in WT and Δ*mreB* strains, including a replicate of the PAMNT054 which was used for all subsequent experiments and its WT comparison. As indicated by the red box, MreC levels were reduced in the Δ*mreB* strains. **(b)** Western blot analysis of FtsZ levels in WT and Δ*mreB* strains. As shown in the red box, FtsZ levels remained unchanged between WT and Δ*mreB* strains.

**Fig. S4**

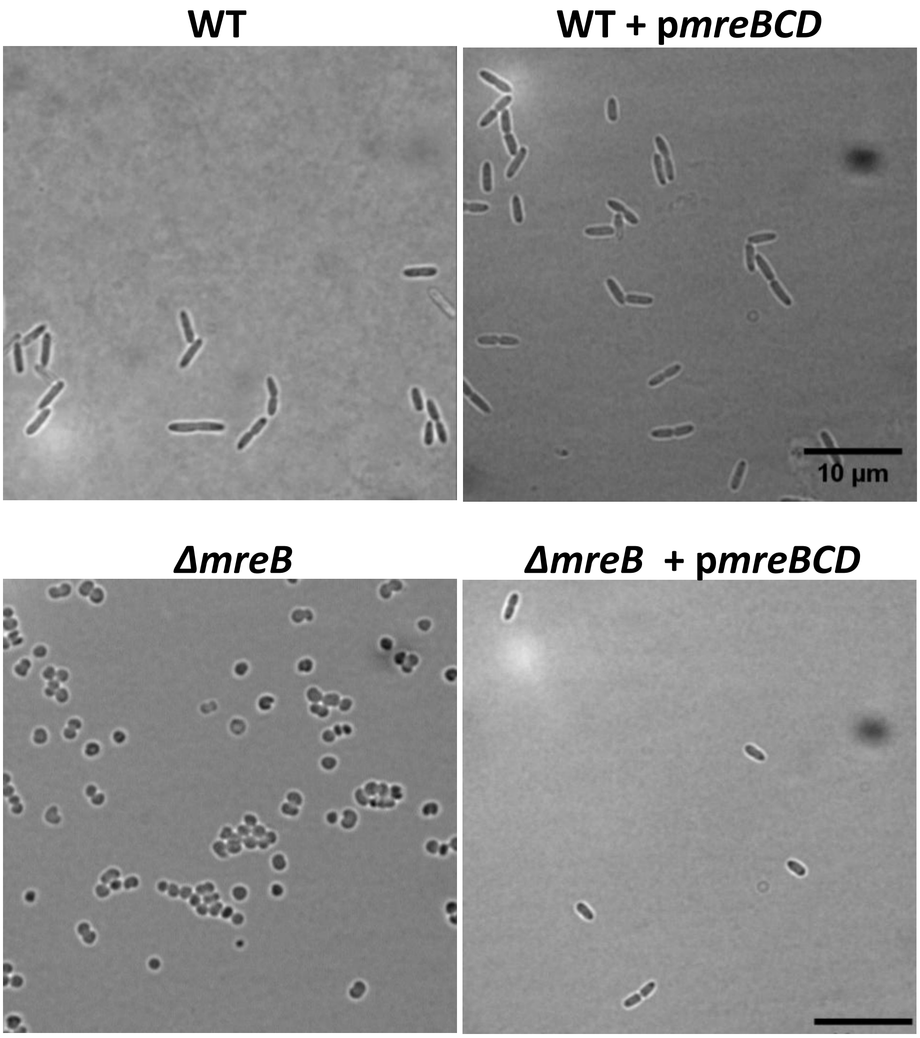


**Fig. S4:** Bright field images of WT, Δ*mreB* with/out an extra copy of ***mreBCD*** under an arabinose-inducible promoter.

**Fig. S5**

**a.**

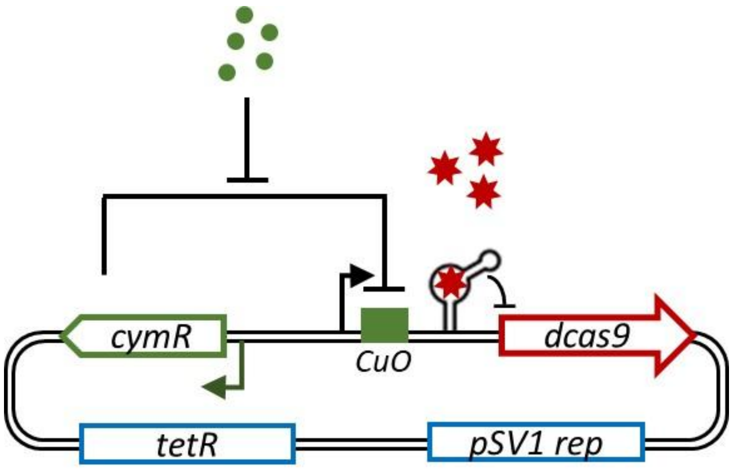


**b.**

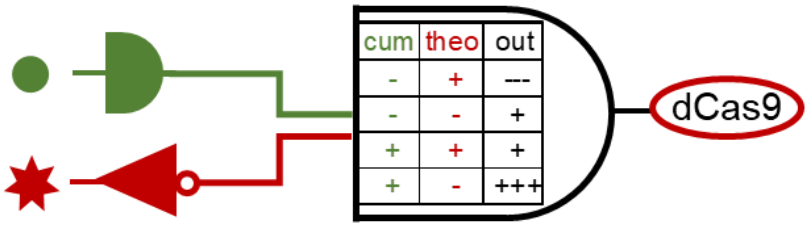


**Fig. S5: (a)** Schematics of the synthetic CRISPRi system used, enabling conditional repression of gene expression in bacteria through two orthogonal small-molecule inputs. Expression of dcas9, encoding a catalytically inactive Cas9 protein for transcriptional repression, is driven by a cumate-inducible promoter repressed by CymR**.** In the absence of cumate (green dots), CymR binds the CuO operator and blocks dcas9 transcription. Upon cumate addition, CymR repression is relieved, leading to induction of dcas9 expression. The system also includes a theophylline-responsive riboswitch (red stars) that controls guide RNA (gRNA) expression. In contrast to typical activator riboswitches, binding of theophylline inhibits gRNA function, thereby blocking CRISPRi activity even in the presence of dcas9 **(b)** Graphical genetic circuit illustrating repression by theophylline and induction by cumate with expected output.

### Fig. S6

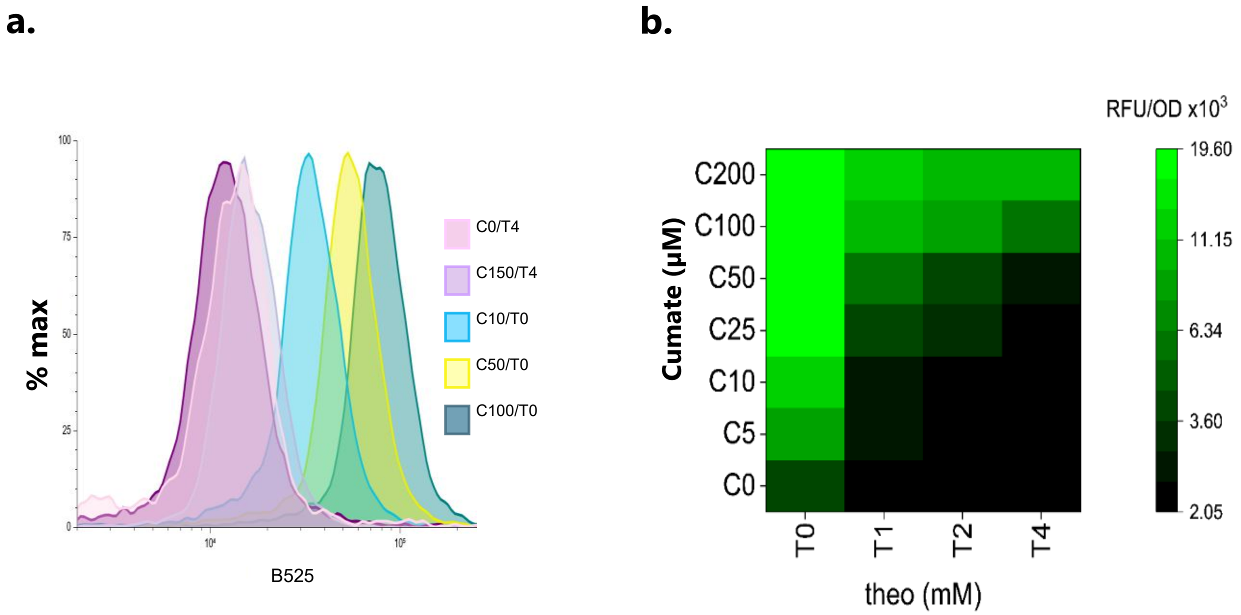


**Fig. S6: (a)** Flow cytometry results of the distribution of green fluorescence intensity (GFP) within the cell population with cumate and theophylline. In each condition, GFP expression was homogeneous throughout the population. **(b)** Different concentrations of cumate and theophylline were tested to determine the optimal dCas9 expression level in a GFP-expressing dCas9 system. dCas9 expression increased with higher cumate concentrations (green), whereas theophylline had a reverse (inhibitory) effect on dCas9 expression (black).

**Fig. S7**

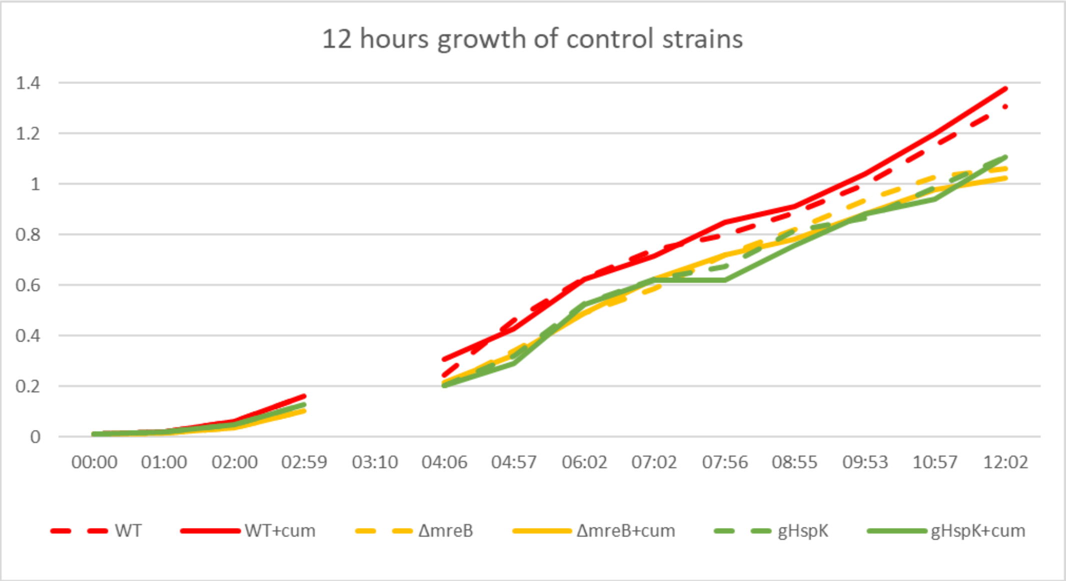


**Fig. S7:** Growth curve of WT, WT + $g^{HspK}$ and ΔmreB strains with and without 200 µM cumate supplementation after 3 hours incubation in LB at 37 °C, 170 rpm. Cumate has no effect on growth rate of control strains at 200 µM concentration.

### Fig. S8

**a.**

| FSC | SSC | Violet SSC | B525 | Y610 | Y690 | Y780 | Treshold | Width |
|---|---|---|---|---|---|---|---|---|
| 500 | 100 | 100 | 500 | 500 | 500 | 500 | SSC-1000_Height | Channel-FSC |

**b.**

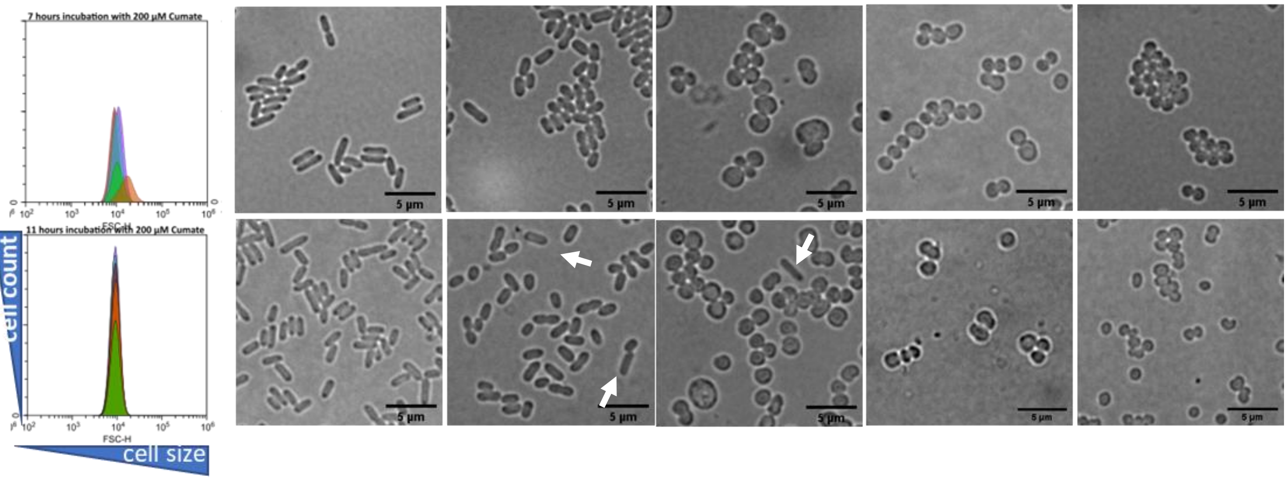


**Fig. S8: (a)** Cytometry acquisition settings. **(b)** Flow cytometry plots and corresponding brightfield (BF) images from the same time points analyzed by flow cytometry. The 7-hour time point includes 5 hours of cumate induction, while the 11-hour time point includes 9 hours of induction. At 5- and 9-hours post-induction of dCas9, WT and *ΔmreB* cells retained their rod-shaped and spherical morphologies, respectively, although both appeared smaller, as expected in the stationary phase when bacterial cells typically reduce their size and metabolic activity. The white arrows indicate rod-shaped cells in the culture. Scale bar is 5 µm.

### Fig. S9

**Fig. S7, 11 hours grown cultures** **Fig. 2d, 12 hours grown cultures**

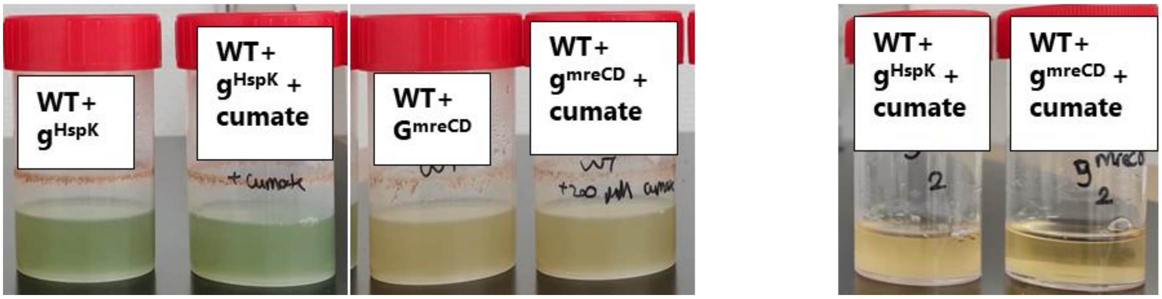


**Fig. S9**: Comparison of **Fig. S8**, 11 hours growth culture with and without 200 µM cumate addition at exponential phase, and **Fig. 2d** starter point of imaging which is after 12 hours of growth with two-time 200 µM cumate addition. WT+ $g^{HspK}$ cultures growth in both conditions while WT + $g^{mreCD}$ did not grow in the second condition (Fig. 2d).

### Fig. S10

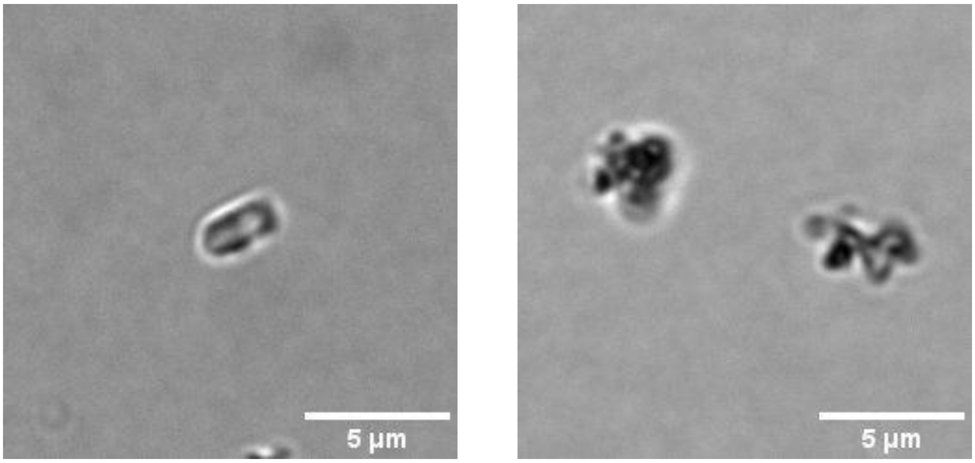


**Fig. S10:** Dead cells from the beginning of the acquisition of *mreBCD* silencing after 12 hours incubation in liquid LB culture supplemented by 2 time 200 µM cumate (first addition at the beginning, second addition after 4 hours incubation at 37°C, 170 rpm). Scale bar is 5 µm.

**Fig. S11**

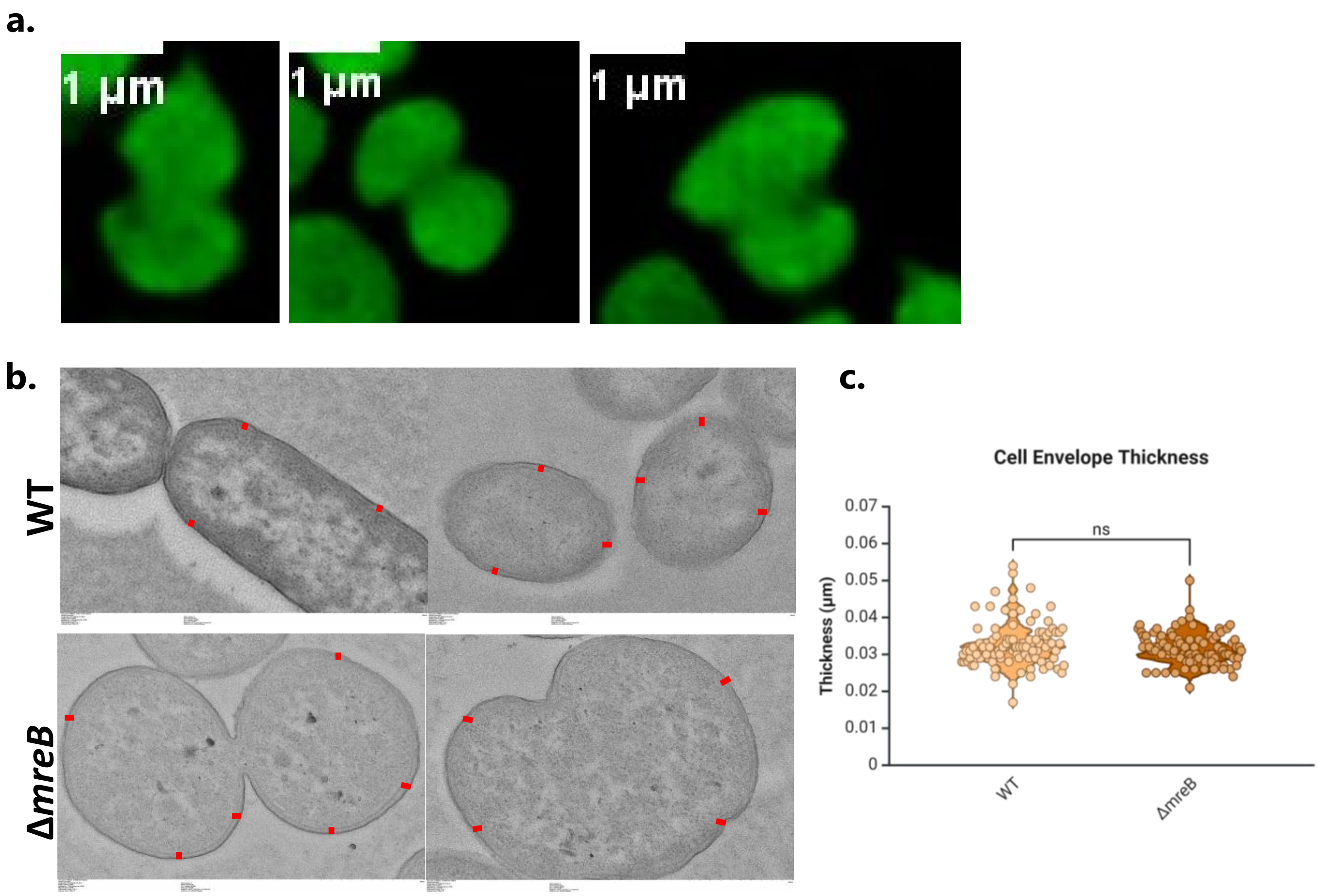


**Fig. S11: (a)** Asymmetrical division of constitutively sfGFP expressed *ΔmreB* cells and deviations from spherical shape were observed, likely due to irregular divisome positioning. Scale bar is 1 µm. **(b)** Transmission Electron Microscopy (TEM) images of sectioned WT and Δ*mreB* samples. Red lines refers measurement area of cell envelope. **(c)** Measurement of cell envelope thickness of WT and Δ*mreB* strains. . Kruskal-Wallis test with Dunn's multiple comparisons test was applied and no significant differences has been found.

**Fig. S12**

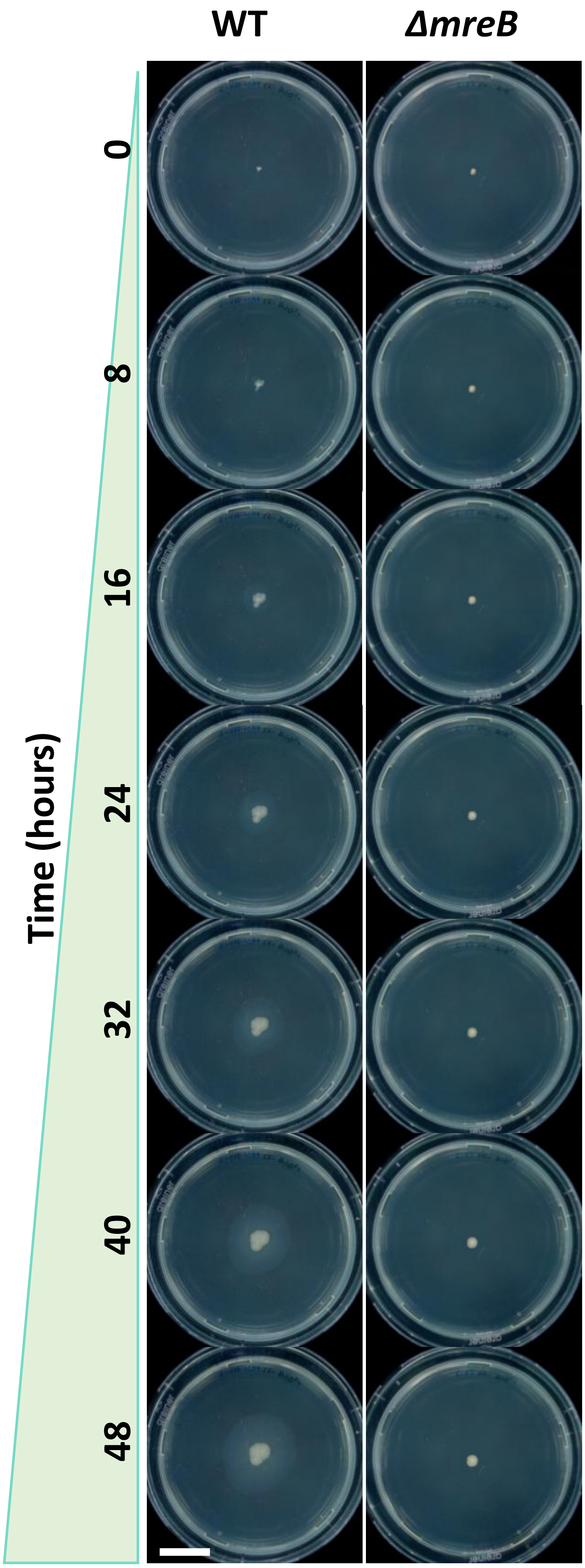


**Fig. S12:** Growth of WT and *ΔmreB* on twitching plate. Growth observed every 8 hour for 2 days at 37°C. Scale bar is 2 cm.

#### Fig. S13

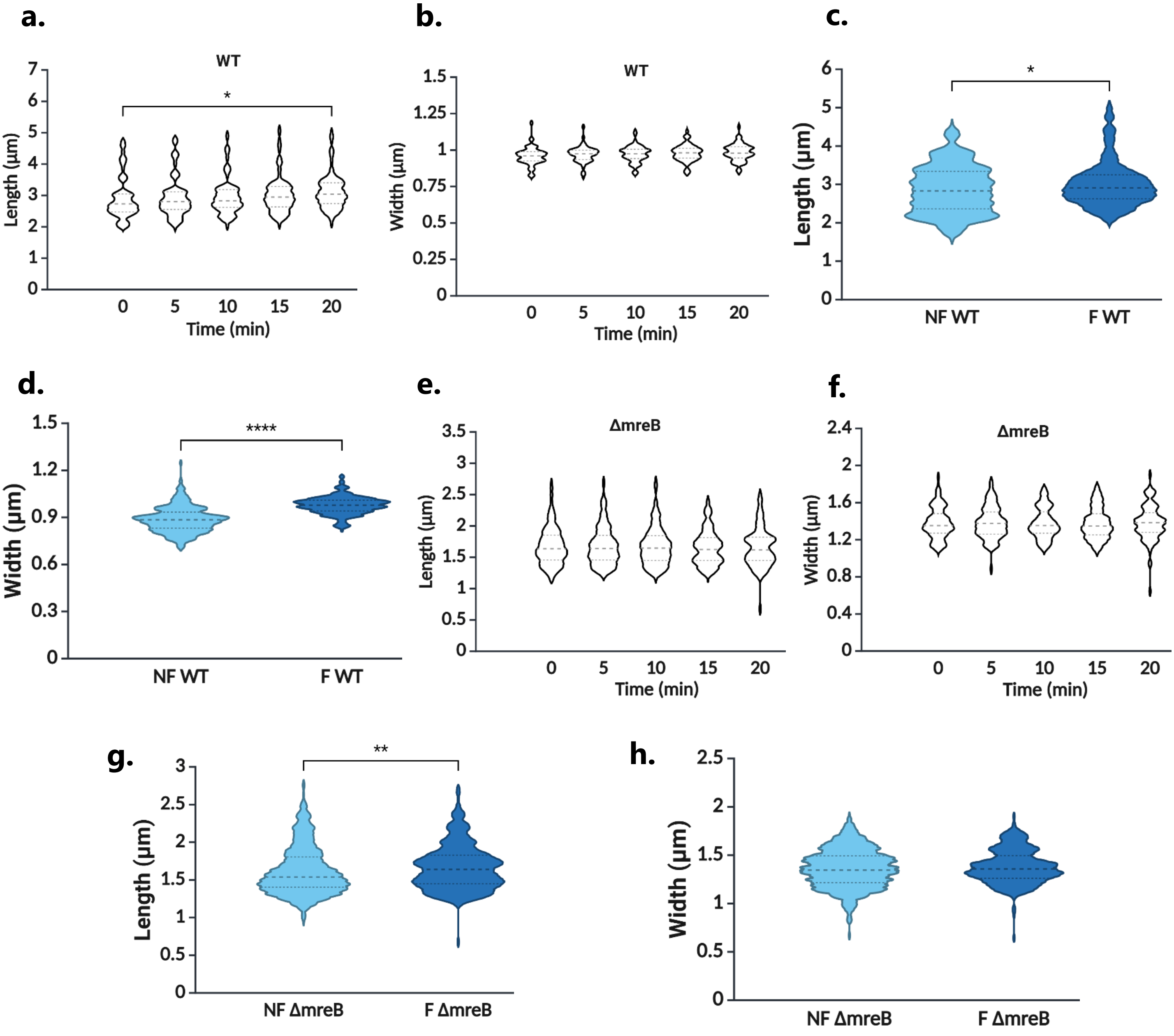


**Fig. S13:** Measurement of Length **(a)** and width **(b)** of WT + mScarletI strain (F WT) in monoculture and its length **(c)** and width **(d)** comparison with non-fluorescent WT (NF WT) strain. Measurement of Length **(e)** and width **(f)** of *ΔmreB* + sfGFP strain (F *ΔmreB*) in monoculture and its length **(g)** and width **(h)** comparison with non-fluorescent *ΔmreB* strain (NF *ΔmreB*). 0 is the first frame after the cells are put on an LB 1% agarose pad. Kruskal-Wallis test with Dunn's multiple comparisons test was applied separately. The initial time point (0) of WT + mScarletI was found to be significantly lower than time point 4 (20 mins) for length, $p = 0.0204$, while the width of WT + mScarletI and both width and length of *ΔmreB* + sfGFP had no significant differences over time. A nonparametric, two-tailed Mann-Whitney U Test was applied for strain length and width comparison separately. In both strains, length is significantly longer than their nonfluorescent types; $p = 0.0171$ for WT + mScarletI and $p = 0.001$ for *ΔmreB* + sfGFP. On the other hand, the width of WT + mScarletI increased ($p < 0.0001$) while it did not change in *ΔmreB* + sfGFP.

### Fig. S14

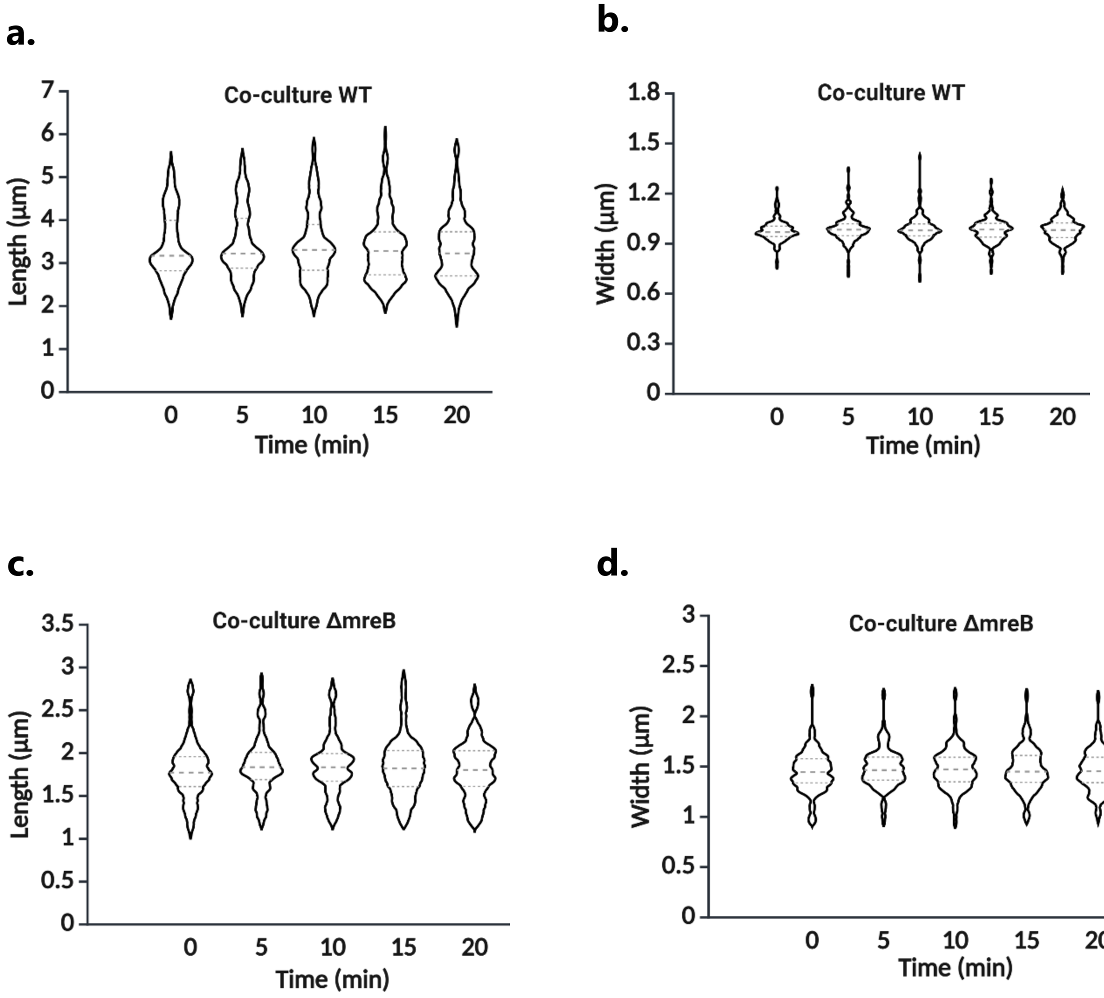


**Fig. S14:** Measurement of Length **(a)** and width **(b)** of WT + mScarletI, length **(c)** and width **(b)** of ΔmreB + sfgfp cells in co-culture over 20 minutes with 5 min intervals. 0 is the first frame after the cells are put on an LB 1% agarose pad. Kruskal-Wallis test with Dunn's multiple comparisons test was applied separately, and no differences were found.

**Fig. S15**

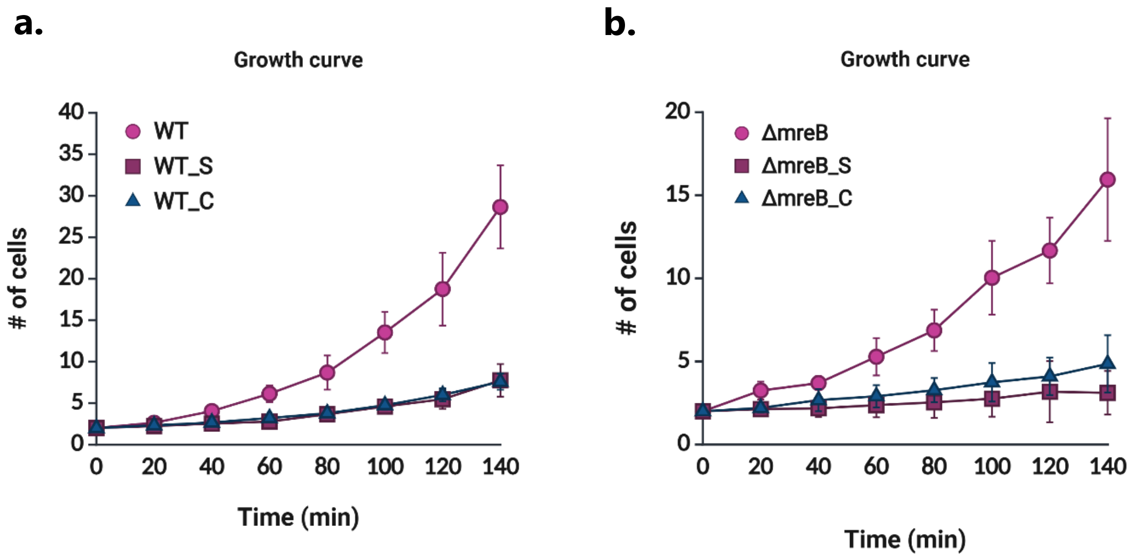


**Fig. S15:** Growth rates of (a) WT and WT + mScarletI in monoculture (WT_S) and co-culture (WT_C), and (b) Δ*mreB* and Δ*mreB* + sfGFP in monoculture (Δ*mreB*_S) and co-culture (Δ*mreB*_C) over 140 mins. No significant differences were observed between monoculture and co-culture conditions for either fluorescent strain. However, expression of fluorescent proteins from plasmids resulted in a reduced growth rate.

#### Fig. S16

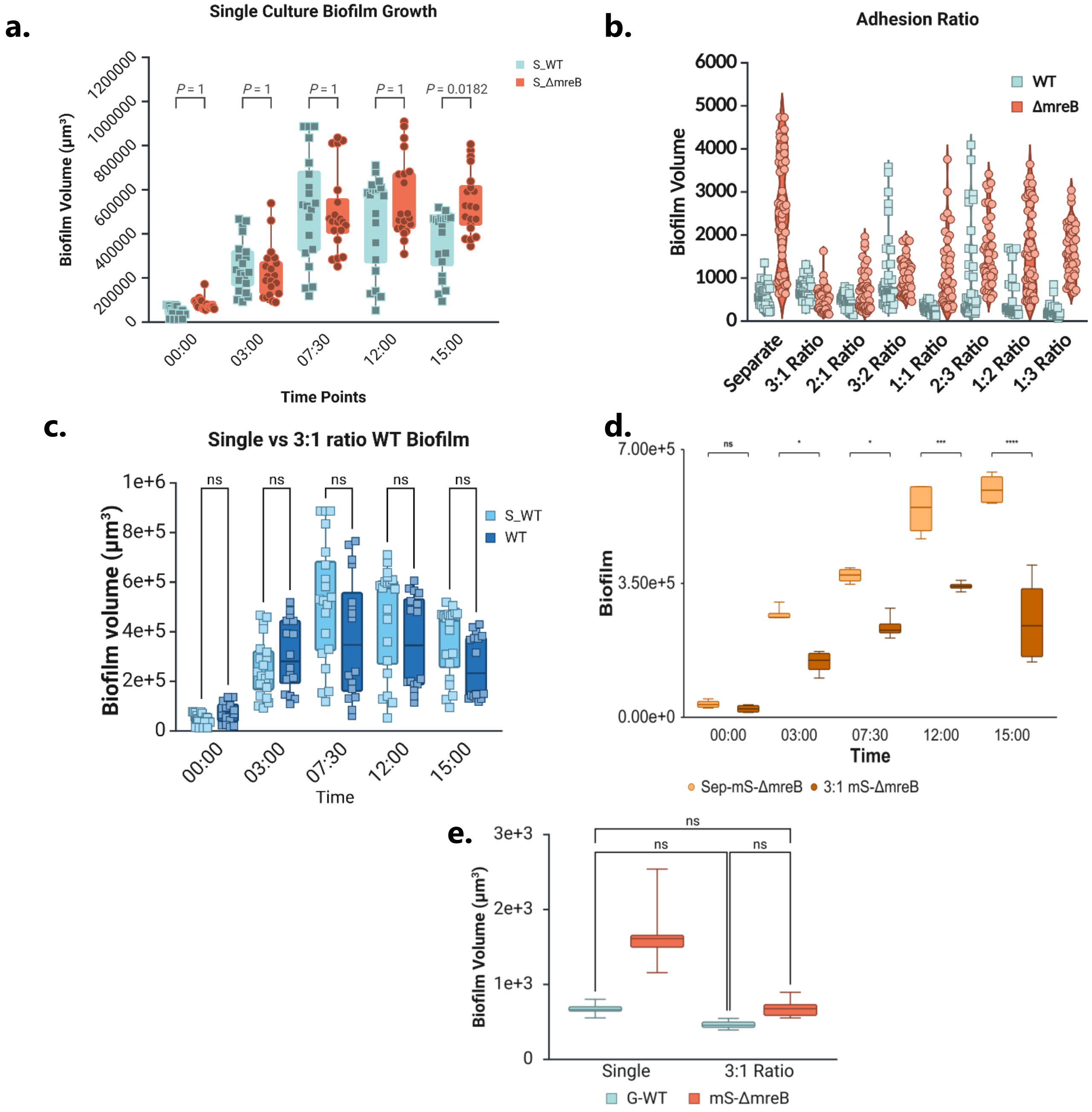


**Fig. S16: (a)** Biofilm volume comparison of WT + mScarletI and Δ*mreB* +sfGFP strains over 15 hours of growth in LB at 37 °C. Kruskal-Wallis test with Dunn's multiple comparisons test was applied, and no significant differences in biofilm volume were observed until the 15-hour time point ($p$=0.0182). **(b)** Comparison of WT + mScarletI and Δ*mreB* +sfGFP ratio at the adhesion step to find the same starter inoculum in co-culture biofilm formation. **(c)** Comparison of biofilm volume of WT + mScarletI in mono and coculture with Δ*mreB* +sfGFP. Kruskal-Wallis test with Dunn's multiple comparisons test was applied, and no significant differences in biofilm volume were observed. **(d)** Comparison of biofilm volume of Δ*mreB* + mScarletI in mono and coculture with WT +sfGFP. TWO WAY ANOVA with Bonferroni test was applied, and significant differences in biofilm volume were observed after 3 hours growth. ($p$=0.005, $p$=0.003, $p$<0.0001, $p$<0.0001 respectively)- **(e)** Quantification of biofilm volume at the adhesion stage (t = 0) for WT + sfGFP and Δ*mreB* + mScarletI strains in monoculture and in co-culture (3:1 ratio, WT:Δ*mreB*). Δ*mreB* exhibits significantly greater initial biofilm volume than WT in monoculture, whereas in co-culture, both strains begin with similar adhesion volumes.

**Fig. S17**

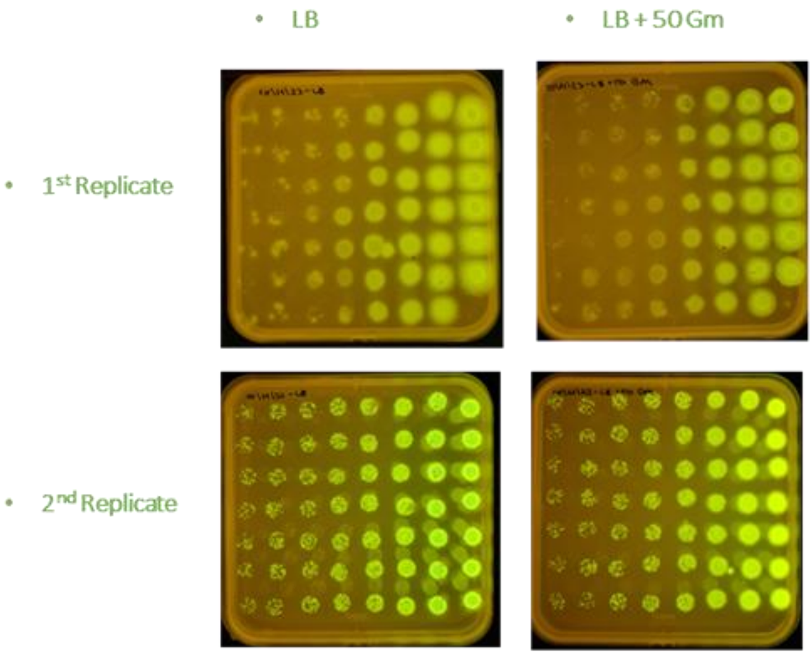


**Fig. S17:** pSEVA627m-sfGFP plasmid expression level in WT with and without 50 mg/L gentamicin. Results indicate that strain did not lose its plasmid over 24 hours.

#### Fig. S18

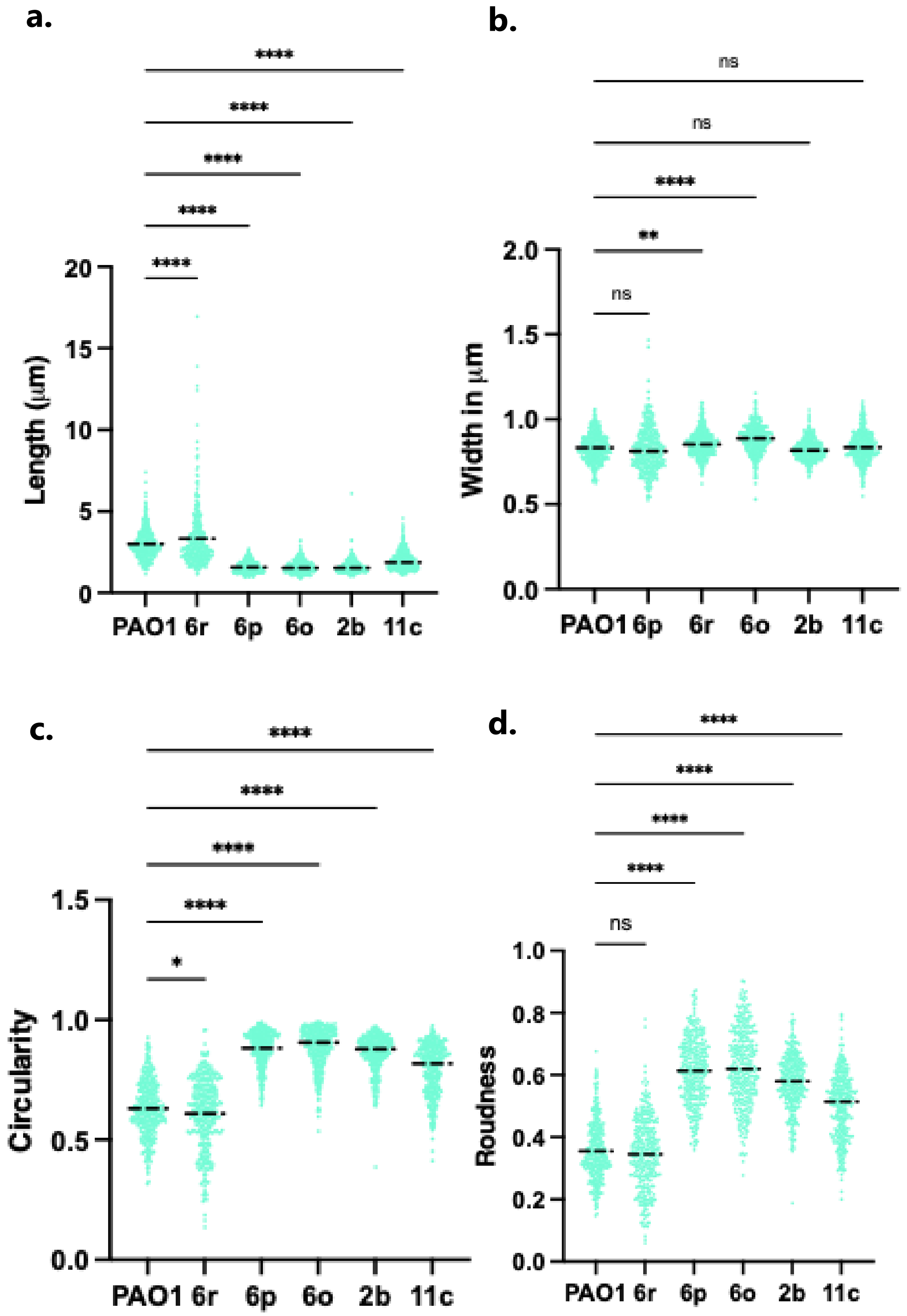


**Fig. S18:** *P. aeruginosa* PAO1 (wild type) and five clinical strains carrying truncated **mreB** sequences were grown in LB rich medium until an $OD_{600}$ of 0.5. Quantitative comparisons of cell **length (a)**, **width (b)**, **circularity (c)**, and **roundness (d)** are shown. The lengths of strains 6p, 6o, 2b, and 11c are significantly shorter than that of the wild-type strain. All strains with reduced length exhibit circularity values close to 1, in contrast to the wild type, which shows a circularity value close to 0.5. Each point refers one cell measurement.
